# Plant plasticity in the face of climate change – CO_2_ offsetting effects to warming and water deficit in wheat. A review

**DOI:** 10.1101/2025.02.10.637370

**Authors:** Meije Gawinowski, Karine Chenu, Jean-Charles Deswarte, Marie Launay, Marie-Odile Bancal

**Author notes:** Corresponding author: Meije Gawinowski, 22 place de l’Agronomie 99120 Palaiseau France. Equally credited senior authors (Marie Launay, Marie-Odile Bancal).

## Abstract

Future crop production will depend on plant plasticity in response to increases in atmospheric CO_2_, mean temperature, heatwave and drought events. The present review intends to highlight the impact of interactions between high CO_2_ levels, warming and water deficit in existing published experimental data in the case of wheat. To do so, we identified experiments quantifying the effects of such interactions on traits related to crop productivity and water use. We used the collected data to estimate plasticity indices assessing compensation and interaction between elevated CO2 and adverse climatic conditions, bringing a new perspective on the matter. In the studied data, even though there is an important variability, we found that crop productivity tends to decrease despite the positive effects of the rise in CO_2_ concentration. Conversely, with elevated CO_2_, water consumption tends to decrease despite the warmer conditions. We hypothesized that the positive effect of CO_2_ on crop productivity is greater under drought conditions, which is confirmed in 54% of the experiments. This review highlights the need to acquire further experimental data under possible future conditions to calibrate and validate crop models: their range of validity requires more thorough testing under the wide range of projected environmental conditions.

## 1. Introduction

### 1.1. Phenotypic plasticity under climate change

Anthropogenic activities increase carbon dioxide (CO_2_) concentration, with a documented rise from 340 to 415 ppm between 1980 and 2020. Future projections range between 540 and 1300 ppm by the end of the century, with a concomitant rise in mean global temperature by up to +4°C (Calvin et al., 2023). This increase in temperature will lead to shifts in climate patterns (IPCC, 2014). Extreme climatic events, such as heatwaves and droughts, are likely to increase in frequency and severity in many cropping areas, posing a growing threat to natural and agricultural ecosystems, particularly when they occur together as combined events (Zscheischler et al., 2018). Plants are sessile organisms; therefore, phenotypic plasticity, defined here as the phenotypic variation in traits in response to the environment (Hallgrimsson et al., 2019), is crucial to their adaptation to varying environmental conditions such as climate change. Plant plasticity to environmental variation assessed by experiments is often limited to the combination of stress modalities and cultural practices. This experimental approach can be expanded by soil-crop modeling (e.g. Chenu et al., 2017; Asseng et al., 2019). However, issues have been raised regarding the accuracy of these models to predict response to future environmental conditions, such as overestimation of CO_2_ effects or under-estimation of combined heat and drought stresses (Chenu et al., 2017; Ahmed et al., 2019). To improve the validity of crop models in the context of climate change, it is hence necessary to better understand the quantitative effects of environmental variables and their interactions. The main objective of this review is to investigate plasticity to such environmental interactions based on published experimental data. Our analysis provides a synthetic overview of the collected data in order to answer three main questions: What are the quantitative impacts on (i) productivity-related traits and (ii) water-related traits from elevated CO_2_ combined with warm temperatures, heatwaves and/or water deficit? (iii) Is the fertilizing effect of CO_2_ stronger under stressed than non-stressed conditions? Plant plasticity to a single environmental factor was widely studied for elevated CO_2_ (eCO_2_) levels (e.g. Ainsworth and Long, 2021), high temperatures (i.e. an increase in mean daily temperature), heatwaves (e.g. Farooq et al., 2011) and water deficit (e.g. Vadez et al., 2024). When plant resources (e.g. CO_2_, water) and growth drivers (e.g. temperature, water) vary outside of their optimal range, they induce stress (e.g. water deficit or excess, heat or cold). A biological stress is defined as a “change in the environment that might reduce or adversely change plant growth or development” (Levitt, 1972; Salisbury and Ross, 1992). Both (i) heat stress, caused by high mean temperatures, hot days, warm nights and/or heatwaves, and (ii) drought stress, caused by soil and/or atmospheric water deficits, are major abiotic stresses known to impair plant growth.

### 1.2. Wheat plasticity in response to key environmental variables

This section describes the separate effects of elevated CO_2_, high temperatures, heatwaves and water deficit with a focus on wheat as one of the major crops delivering calories worldwide (Cassidy et al., 2013). Stress impacts on key processes are summarized in Figure 1.

**Figure 1.**
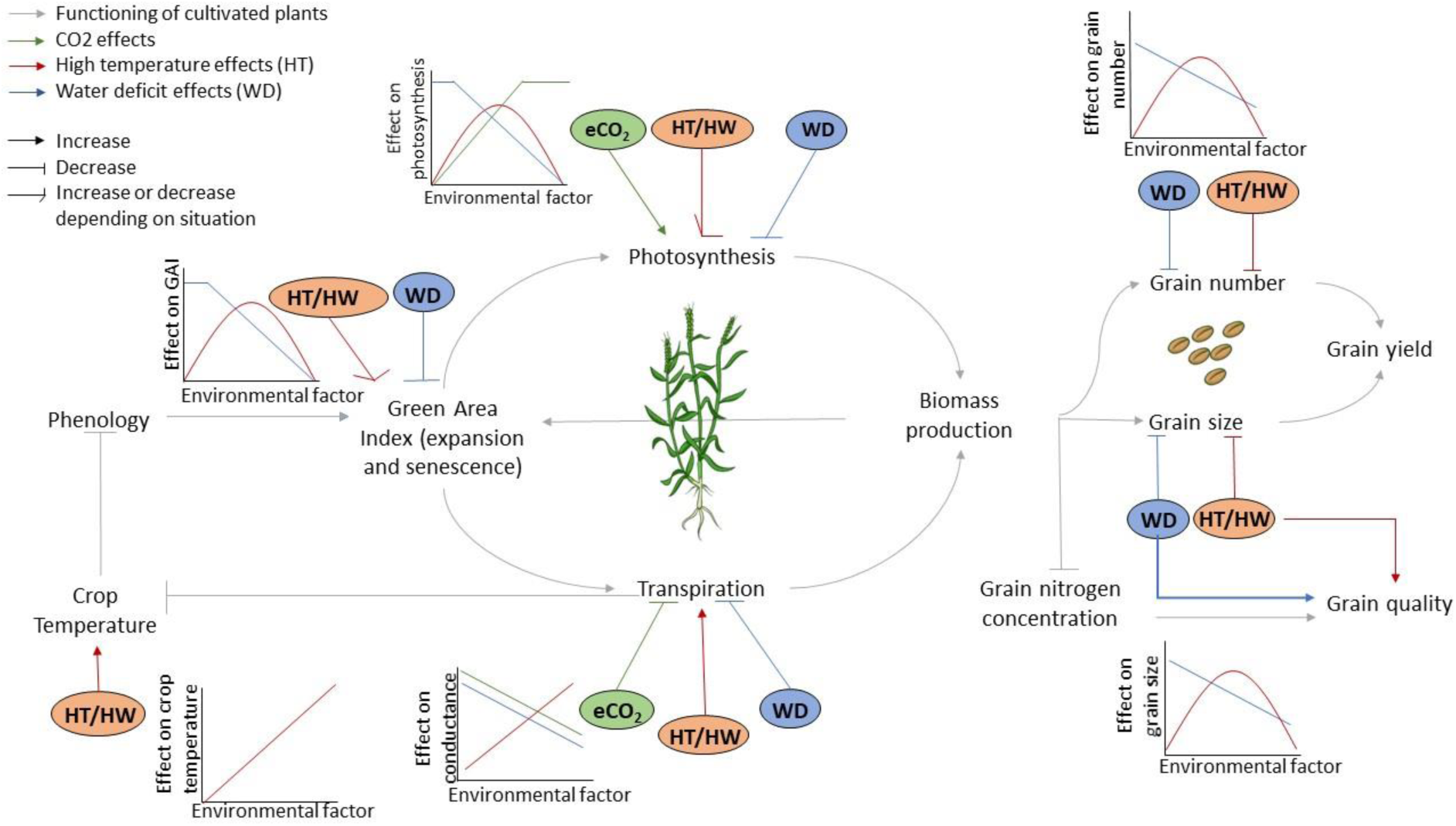
Conceptual model of wheat plasticity showing the different shapes of response curves to single environmental factors, i.e. elevated CO_2_ (eCO_2_, green, see subsection 1.2.1), warming (red, see subsection 1.2.2) through high temperatures (HT) or heatwaves (HW) and water deficit (WD, blue, see subsection 1.2.3). For most processes, the effect of warming depends on the proximity to optimal temperatures. The two key processes impacted by the three climatic factors are photosynthesis (increased by eCO_2_, decreased by WD and increased or decreased by warming depending on temperature) and transpiration (decreased by eCO_2_ and WD, increased by warming). Warming and water deficit also directly impact green area index, grain number, grain size and grain quality, that are also indirectly impacted by eCO_2_. Warming also strongly affects phenology by shortening crop duration.

#### 1.2.1. Wheat plasticity to elevated CO2

Photosynthesis and stomatal conductance are the two main processes displaying plasticity in response to eCO_2_ (Figure 1). Unlike in C4 plants, Rubisco in C3 plants is not saturated in CO_2_ (Long et al., 2004), so photosynthesis can be stimulated by elevated atmospheric CO_2_ concentration (eCO_2_). Hence in the absence of stress, eCO_2_ in C3 plants increases biomass, tillering, leaf area, yield components and ultimately yield (Figure 1; Long et al., 2004; Kadam et al., 2014). However, eCO_2_ does not necessarily enhance N uptake, so increases in biomass accumulation under eCO_2_ result in nitrogen dilution (Högy and Fangmeier, 2008) and lower concentrations of nitrogen and protein in grains (Figure 1; Kadam et al., 2014). An increase in CO_2_ concentration also greatly decreases stomatal conductance and plant transpiration, improving water-use-efficiency (WUE) in crops like wheat (Li et al., 2023). Since the 1980s, the effects of eCO_2_ have been studied in pots or in the field, in diverse experimental conditions, including controlled growth chambers, greenhouses, closed-top chambers (CTC), open-top chambers (OTC) and free air CO_2_ enrichment experiments (FACE). Enclosure studies (chamber, CTC, OTC) are implemented on a smaller scale and even if some are partially open to the atmosphere, they are not representative of fully open-air conditions compared to FACE studies. These experiments have focused on different biological levels (e.g. organ, plant and canopy levels) and different time scales (e.g. minute, hour, day, crop cycle). In wheat, the results of such experiments were summarized and evaluated in various meta-analyses and reviews (Ainsworth and Long, 2021, 2005). Overall, increasing CO_2_ concentration from 380 to 550 ppm in non-stressful conditions resulted in an average yield gain of +31% in enclosed studies and +13% in FACE studies (Long et al., 2006). Under elevated CO_2_ (546-586 ppm) levels, grain protein concentration decreased by 6.3% on average compared to ambient CO_2_ across seven FACE experiments at different locations (Myers et al., 2014). For water balance, a 22% decrease in stomatal conductance was reported for an increase in CO_2_ concentration from 366 to 567 ppm in C3 species (Ainsworth and Rogers 2007). Even for the effect of CO_2_ alone, it is difficult to obtain clear consistent response curves due to the highly diverse conditions used in experimental set up, and the impact that other environmental factors have on crops (Ainsworth and Long, 2021).

#### 1.2.2. Plasticity to warming

In this study, we consider two types of temperature effects related to warming: (i) the homogeneous rise in mean temperatures, referred to here as ‘high temperatures’, and (ii) the occurrence of extreme heat stress events, referred to here as ‘heatwaves’, which frequency, duration and intensity are predicted to increase with climate change. Thermal stress results from non-optimal temperatures due to either warming or cold/frost (Barlow et al., 2015). This review only focuses on warming, as this is the principal effect of current climate change (Collins and Chenu, 2021), but increasing average temperatures also increases frost damage in some regions by advancing phenology (Bai et al., 2022; Zheng et al., 2015). A homogeneous increase in mean temperatures up to an optimum might increase photosynthetic activity (Figure 1) but can also generate a stress when process-dependent thresholds are exceeded, notably around flowering and grain filling. Heatwaves harm developing plants at all stages by damaging the photosynthetic machinery, reducing tillering, altering pollen, and causing kernel abortion and physiological injuries (Hunt et al., 2018; Parthasarathi et al., 2022). Increases in temperature typically result in a shorter growth period (Girousse et al., 2021; Zheng et al., 2012), limiting resource acquisition (e.g. light, N, CO_2_) and biomass production. Heatwaves also cause major yield losses through the disruption of meiosis in pollen, grain abortion, accelerated senescence, lower levels of grain filling, and a loss of assimilate remobilization (Dolferus, 2014; Yahya et al., 2022). Sub-optimal temperatures also affect grain quality by increasing grain protein content and altering starch composition (Kadam et al., 2014; Triboi et al., 2006). If heatwaves are more studied in terms of daytime temperatures, warm nights also have an important impact by reducing spikelet fertility, decreasing grain number and size (Farooq et al., 2011; Prasad et al., 2008). Transpiration is increased under high temperatures as a common cooling strategy which reduces leaf temperature and mitigates the negative effect of heat (Farooq et al., 2011). Hence, temperature affects diverse processes displaying different levels of plasticity during both the vegetative and reproductive phases of plant growth. Temperature effects have been studied in various conditions, in growth cabinets/rooms, greenhouses, heat chambers placed over crops in the field, temperature gradient tunnels and T-FACE (FACE facilities with infrared heaters for temperature variation). However, the effects of temperature can also be studied by modifying sowing date and analyzing the effect of naturally occurring heatwaves, e.g. with heat chambers (Thistlethwaite et al., 2020) or the use of a photoperiod-extension method to synchronize the phenology of plants/genotypes (Ullah et al., 2023, 2024). Temperature effects have also been analyzed with historical yield and climate data or simulation studies. Such analyses concluded to a decrease in global wheat yield in response to an increase in global mean temperature of +1°C: −6% in Asseng et al. (2015) with multi-model simulations, −4.1 to −6.4% in Liu et al. (2016) with point or grid based models and regressions on historic data, −6.9% in Ottman et al. (2012) in a field study, −6% in Zhao et al. (2017) from global grid or local point based models, statistical regressions and field-warming experiments. However, the magnitude of this decrease is also dependent on the location, for example the simulation study conducted by Asseng et al. (2011) with APSIM-NWHEAT concluded to a decrease in Australian wheat yields by up to 50% for temperatures of +2°C, mostly attributed to temperatures above 34°C that impacted to increased leaf senescence and grain filling, notably with an additional indirect effect through reduced soil moisture at flowering. However, it is more difficult to generalize the effects of occasional thermal stress like heatwaves (Ababaei and Chenu, 2020; Lobell et al., 2015; Ullah et al., 2023b). Plastic response to heat varies between genotypes (Ullah et al., 2023a) as breeding for heat tolerance and avoidance is already performed. Adaptation strategies are also considered, notably with the use of earlier cultivars and/or earlier sowing dates to avoid major stress periods (Collins and Chenu, 2021; Gouache et al., 2012).

#### 1.2.3. Plasticity to water deficit

Water deficit, arguably the most important abiotic constraint on crop productivity, is defined as a shortage of atmospheric and/or soil water resulting in substantial morphological, biochemical, physiological and molecular changes (Sallam et al., 2019; Vadez et al., 2024). During the vegetative period, water deficit decreases leaf elongation and photosynthetic activity (Leveau et al., 2021; Prasad et al., 2011) and increases pollen sterility by impairing meiosis (Ji et al., 2010) thus leading to lower rates of grain set (Figure 1). Water deficits during grain filling, which are frequent in many regions (Chenu et al., 2011, 2013a; Chenu, 2015), decrease grain size by altering photosynthesis and shortening the grain-filling period through an acceleration of senescence (Christopher et al., 2016, 2014; Farooq et al., 2009). Water deficit can also impair grain quality with reduced grain protein content (Wan et al., 2022) and impaired quality of flour proteins (Yang et al., 2023). Finally, water deficit decreases stomatal conductance and transpiration, thereby increasing water-use efficiency (Ahmad et al., 2018; Chenu et al., 2018; Collins et al., 2021; Qaseem et al., 2019). Drought stress may result from a deficit of soil and/or atmosphere water through a lack of precipitations, low soil moisture and/or high vapor pressure deficit, but their respective effects on plant growth are still misunderstood (Aguirre et al., 2021). Indeed, most studies to date ignore the role of atmospheric water deficit and have only investigated drought stress by decreasing soil water content, either with controlled irrigation (Schoppach and Sadok, 2012; Shazadi et al., 2024), by removing rainfall with rainout shelters (Hoover et al., 2018) or by adjusting the water added by irrigation in arid environments (Wen et al., 2023), especially for studies combining water deficit with eCO_2_ and/or warming. Plant water relations are greatly impacted by the approach for water restriction (Puértolas et al., 2017), as well as growth conditions in pots or field (Ogbaga et al., 2020). Yield losses due to water deficit vary considerably, depending on the duration and timing of the stress, but also genotype and environmental conditions (Farooq et al., 2009; Prasad et al., 2011; Qaseem et al., 2019). Cultivars display variability in response to water deficit due to breeding for drought tolerance and avoidance, notably through earliness, efficiency of water uptake, transpiration efficiency, and stay-green traits (Farooq et al., 2014; Rebetzke et al., 2009).

### 1.3. Baseline hypotheses and methodology of this review

High CO_2_ levels and increases in mean temperatures, heatwaves and water deficit have direct effects on many physiological processes, including two fundamental ones: photosynthesis and stomatal conductance (Figure 1). However, when considered individually, these environmental variables may have opposite effects on plants. Overall, plants will respond to the combinations of all environmental factors, including complex trait-trait and trait-environment interactions (Slafer et al., 2023; Vadez et al., 2024). The objective of this review is to summarize and unravel plant responses to combinations of eCO_2_, high temperatures, heatwaves and/or water deficit in crops, focusing on wheat. We used published experimental data to investigate wheat plasticity under such combinations. We focused on (i) productivity-related traits (PRTs), such as yield, plant biomass and photosynthesis per unit area and (ii) water-related traits (WRTs), such as stomatal conductance, transpiration and water use efficiency (WUE). These traits are measured at different scales depending on the studies. The articles included in this analysis are listed and detailed in Tables S1 to S5. They were identified by searching Web of Science for the following keywords in titles: “wheat”, “CO_2_”, “temperature”, “heat”, “drought”, “water”, and from the meta-analysis performed by Zhu et al. (2023). Studies that did not provide exploitable data were excluded. In each study, mean values of productivity and water-related traits were retrieved to compute relative trait variations. Observations specific to an experiment, a stress intensity, a year and a cultivar were averaged over replicates. The extracted data are freely available from the INRAE space of the *Recherche Data Gouv* repository https://doi.org/10.57745/ZEJIEW. The studied experimental data was used to investigate three questions regarding wheat plasticity in response to interactions between the effects of high CO_2_ levels, increases in temperature and water deficit referred to as “future-like” conditions (Figure 2):

- Focus 1: the variation in a given trait *T* between (i) current non-limiting conditions, i.e. ambient CO_2_, well-watered conditions and ambient temperature, and (ii) the future-like conditions under eCO_2_, i.e. high temperature, heatwaves, and/or water deficit (WD) combined with eCO_2_. This variation should provide information about the ability of CO_2_ increase to compensate for the effects of heat and/or water deficit. It was estimated by calculating a plasticity index Δ _[aCO2_ × _noS] vs [eCO2_ × _S]_(T) defined as the relative difference (%) in the considered trait T between (i) ambient atmospheric CO_2_ (aCO_2_) and no stress (noS; T _[aCO2_ × _noS]_) and (ii) ‘future-like’ elevated CO_2_ (eCO2) with a stress (S; T _[eCO2_ × _S]_):

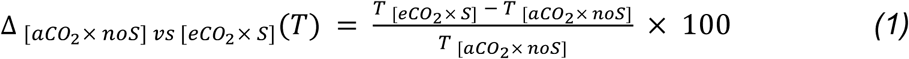

This focus is declined for productivity-related traits (PRTs, such as yield, biomass or photosynthesis, F1.1) and for water-related traits (WRTs, such as stomatal conductance, transpiration or WUE, F1.2).

**Figure 2.**
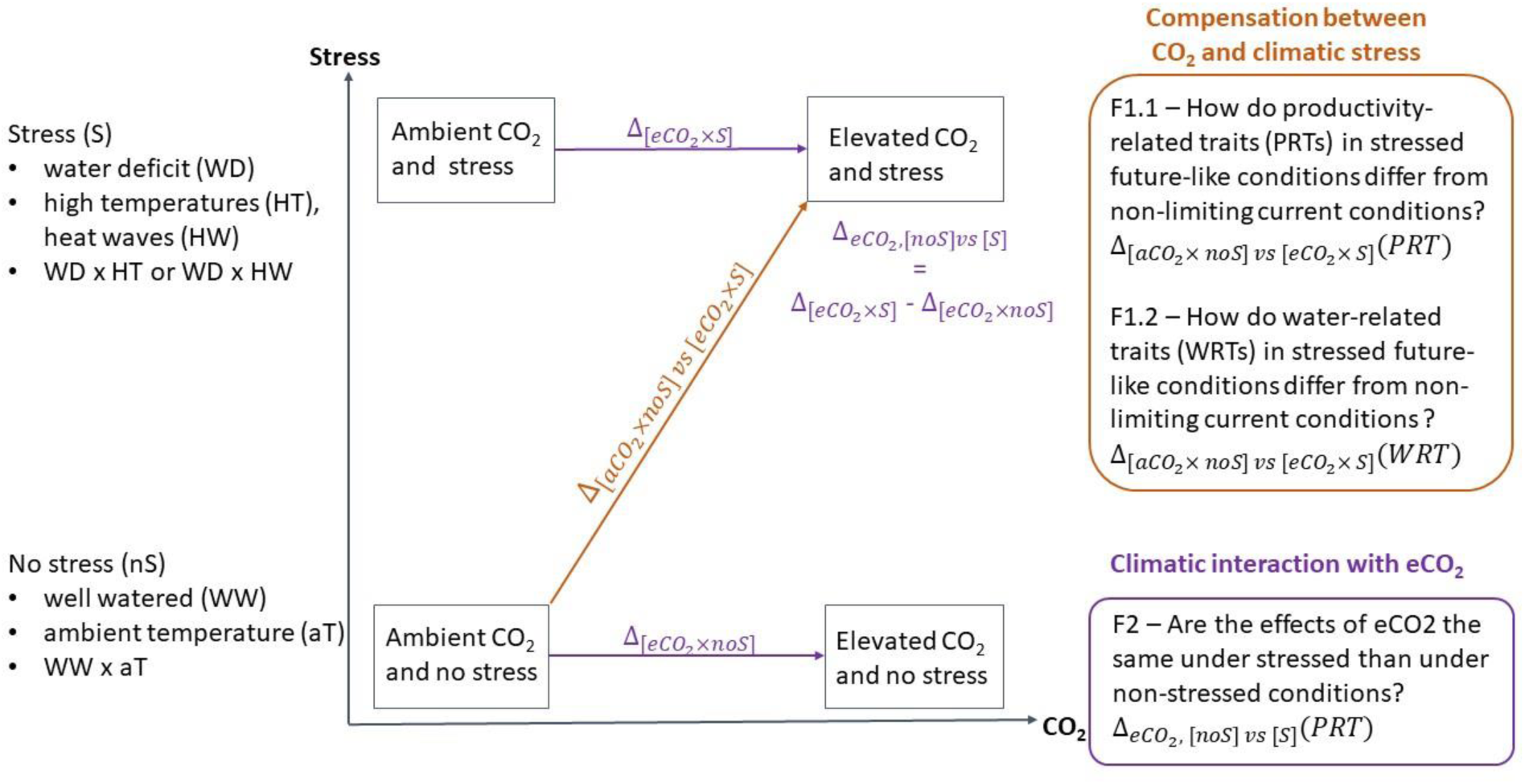
Main focuses of the review and definition of key plasticity indices. Brown arrows represent focus F1 investigating the evolution of productivity-related traits (PRTs, F1.1) with Δ_[aCO2_ × _noS] vs [eCO2_ × _S]_ (PRT), and water-related traits (WRTs, F1.2) with Δ_[aCO2_ × _noS] vs [eCO2_ × _S]_ (WRT) between non-limiting current conditions and future-like conditions. Purple arrows represent focus F2 investigating the interaction between eCO2 and stress on productivity-related traits (F2) with Δ_eCO2, [noS] vs [S]_ (PRT).

- Focus 2 (F2): the difference in the effect of eCO_2_ on crop productivity-related traits (PRT) in non-stressed (noS) and stressed (S) conditions, which should reveal interactions between eCO_2_ and stress factors. The following index was calculated to quantify the variation in F2:

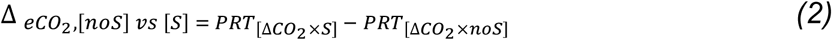

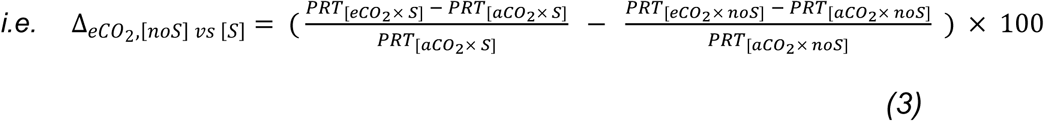

These three questions were studied for different types of climatic interactions:

- Combinations of eCO_2_ with water deficit (WD, Figure 3)

- F1.1 “How do productivity-related traits (PRTs) in stressed future-like conditions with water deficit (WD) under eCO_2_ differ from non-limiting current conditions?” Δ _[aCO2_ × _WW] vs [eCO2_ × _WD]_ (PRT)
- F1.2 “How do water-related traits (WRTs) in stressed future-like conditions with water deficit (WD) under eCO_2_ differ from non-limiting current conditions?” Δ _[aCO2_ × _WW] vs [eCO2_ × _WD]_ (WRT)
- F2 “Are the effects of eCO_2_ on production-related traits the same under stressed (water deficit, WD) and non-stressed conditions (well-watered, WW)?” Δ _CO2, [WW] vs [WD]_ (PRT)
- Combinations of eCO_2_ with warming through high temperatures (HT) or heatwaves (HW)

- F1.1 “How do productivity-related traits (PRTs) in stressed future-like conditions with warming (HT or HW) under eCO_2_ differ from non-limiting current conditions?” Δ _[aCO2_ × _aT] vs [eCO2_ × _HT]_ (PRT) (Figure 4) and Δ _[aCO2_ × _aT] vs [eCO2_ × _HW]_ (PRT) (Figure 5)
- F1.2 “How do water-related traits (WRTs) in stressed future-like conditions with warming (HT or HW) under eCO_2_ differ from non-limiting current conditions?” Δ _[aCO2_ × _aT] vs [eCO2_ × _HT]_ (WRT) (Figure S3) and Δ _[aCO2_ × _aT] vs [eCO2_ × _HW]_ (WRT) (Figure 5)
- F2 “Are the effects of eCO_2_ on production-related traits the same under stressed (warming, HT or HW) and non-stressed conditions (ambient temperature, aT)?” Δ _eCO2, [aT] vs [HT]_ (PRT) (Figure S4A) and Δ _CO2, [aT] vs [HW]_ (PRT) (Figure S4B)
- Combinations of eCO_2_ with water deficit and warming through high temperatures (Figure 6) or heatwaves (Figure 7)

- F1.1 “How do productivity-related traits (PRTs) in stressed future-like conditions with water deficit (WD) and warming (HT or HW) under eCO_2_ differ from non-limiting current conditions?” Δ _[aCO2_ × _WW_ × _aT] vs [eCO2_ × _WD_ × _HT]_ (PRT) and Δ _[aCO2_ × _WW_ × _aT] vs [eCO2_ × _WD_ × _HW]_ (PRT)
- F1.2 “How do water-related traits (WRTs) in stressed future-like conditions with water deficit (WD) and warming (HT or HW) under eCO_2_ differ from non-limiting current conditions?” Δ _[aCO2_ × _WW_ × _aT] vs [eCO2_ × _WD_ × _HT]_ (WRT) and Δ _[aCO2_ × _WW_ × _aT] vs [eCO2_ × _WD_ × _HW]_ (WRT)

**Figure 3.**
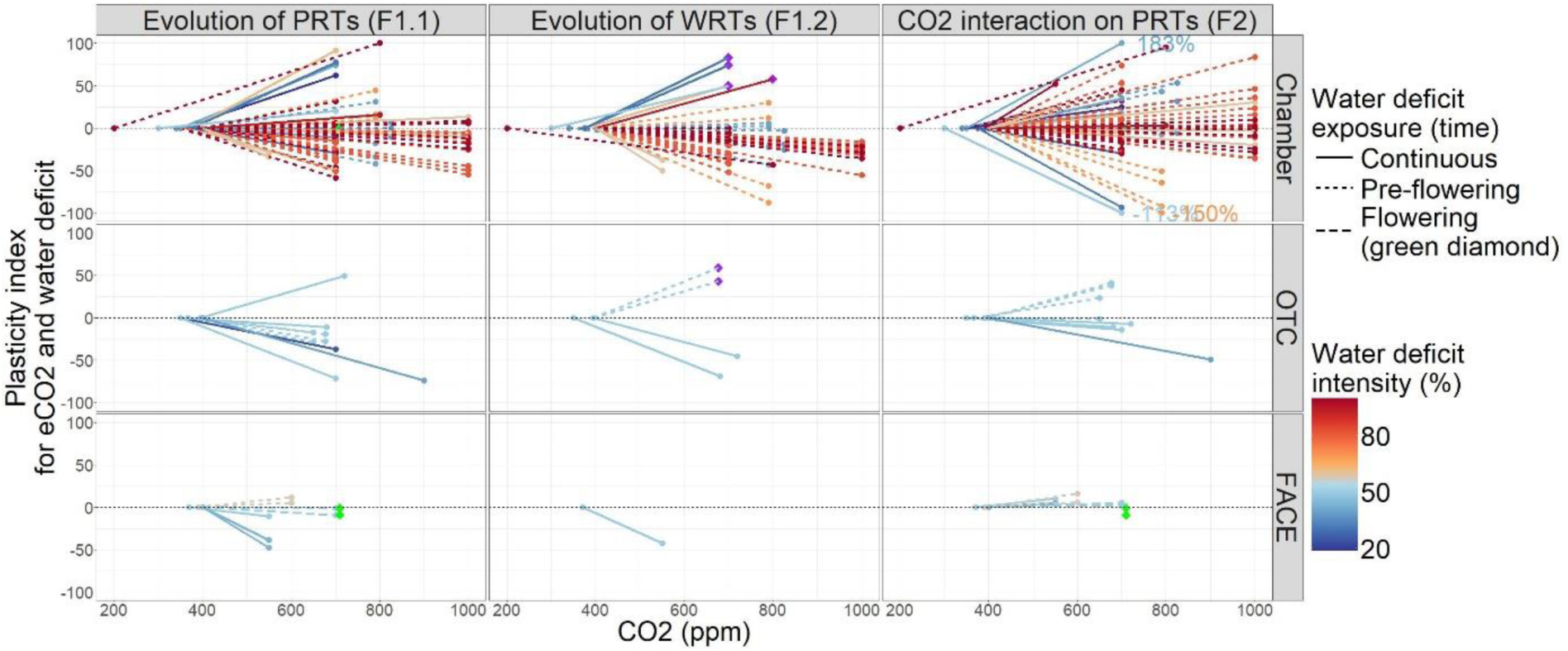
Relative reaction norms for the evolution of productivity (F1.1, Δ_[aCO2_×_WW] vs [eCO2_×_WD]_ (PRT), Eq 1) and water-related traits (F1.2, Δ_[aCO2_×_WW] vs [eCO2_×_WD]_ (WRT), Eq 1) as well as CO_2_ interaction with water deficit on productivity-related traits (F2, Δ _eCO2, [WW] vs [WD]_ (PRT), Eq 2). Productivity traits include biomass and yield variables, water-related traits include stomatal conductance, transpiration and WUE (purple diamonds) variables. Results are presented for different experimental facilities, i.e. chambers, open-top chambers (OTCs) and free-air CO_2_ enrichment (FACE). Colors represent different water deficit intensities (ratio of water inputs between treatments, in %) and line types represent different onset timings for drought stress. The black dashed horizontal lines indicate 0%. Display is limited between −100 and +100% so values beyond are limited to the bounds and the actual values are labelled.

**Figure 4.**
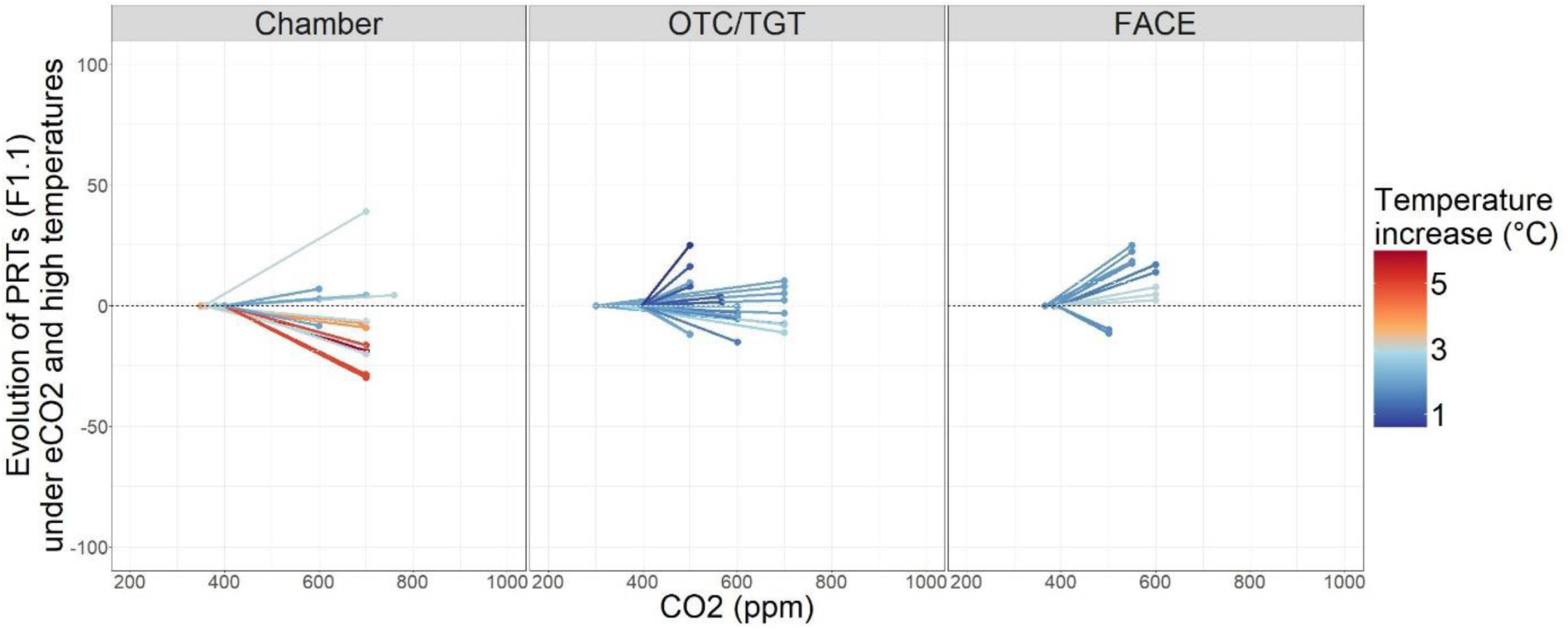
Relative reaction norms for the evolution of productivity (F1.1, Δ_[aCO2_×_aT] vs [eCO2_× *HT*_]_ (PRT), Eq 1) under combined eCO_2_ and high temperatures. Productivity traits include biomass and yield variables. Results are presented for different experimental facilities, i.e. chambers, open-top chambers (OTCs) and free-air CO_2_ enrichment (FACE). Colors represent different temperature increment. The black dashed horizontal lines indicate 0%.

**Figure 5.**
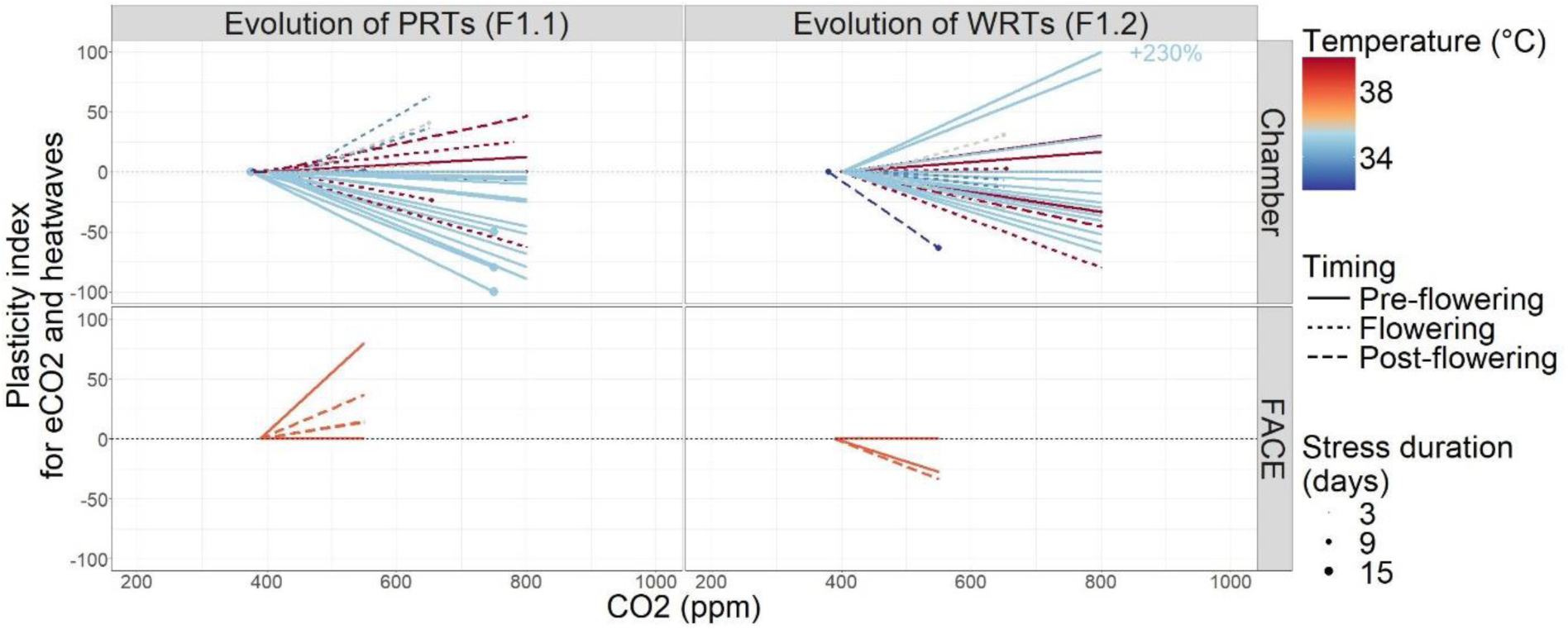
Relative reaction norms for the evolution of productivity (F1.1, Δ_[aCO2_×_aT] vs [eCO2_× *H*_W]_ (PRT), Eq 1) and water-related traits (F1.2, Δ_[aCO2_×_aT] vs [eCO2_×_HW]_ (WRT), Eq 1) for eCO_2_ and heatwaves. Productivity traits include photosynthesis and yield variables, water-related traits include stomatal conductance variables. Results are presented for different experimental facilities, i.e. chambers, open-top chambers (OTCs) and free-air CO_2_ enrichment (FACE). Colors represent heatwave temperatures, line types represent different onset timings for heatwaves and point sizes represent the duration of heatwaves. The black dashed horizontal lines indicate 0%. Display is limited between −100 and +100% so values beyond are limited to the bounds and the actual values are labelled.

**Figure 6.**
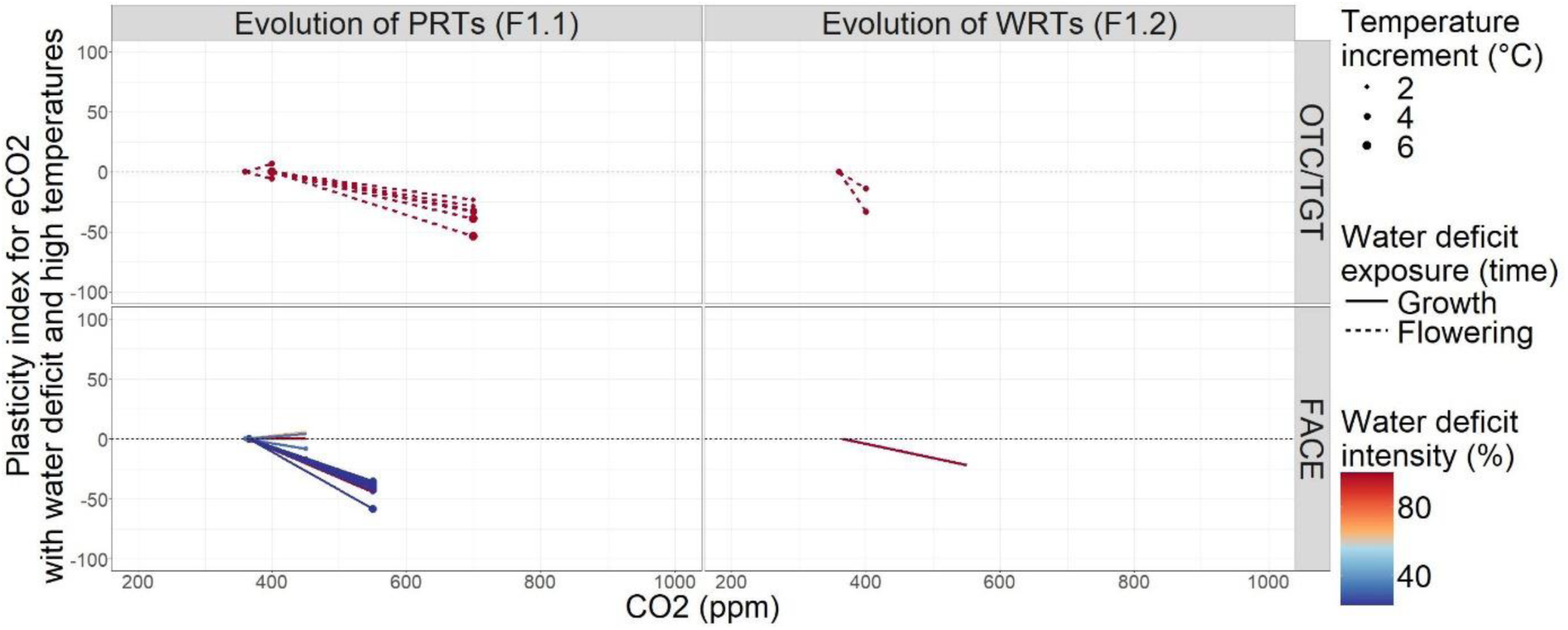
Relative reaction norms for the evolution of productivity (F1.1, Δ_[aCO2_×_WW_× *aT*_] vs [eCO2_×_WD_×HT_]_ (PRT), Eq 1) and water-related traits (F1.2, Δ_[aCO2_×_WW_× *aT*_] vs [eCO2_×_WD_×HT_]_ (WRT), Eq 1) for eCO_2_, water deficit and high temperatures. Productivity traits only include yield variables, water-related traits only include stomatal conductance variables. Results are presented for different experimental facilities, i.e. open-top chambers and temperature gradient tunnels (OTC/TGT) and free-air CO_2_ enrichment (FACE). Colors represent different water deficit intensities (ratio of water inputs between treatments, in %), point sizes represent different temperature increments and line types represent the onset of stress.

**Figure 7.**
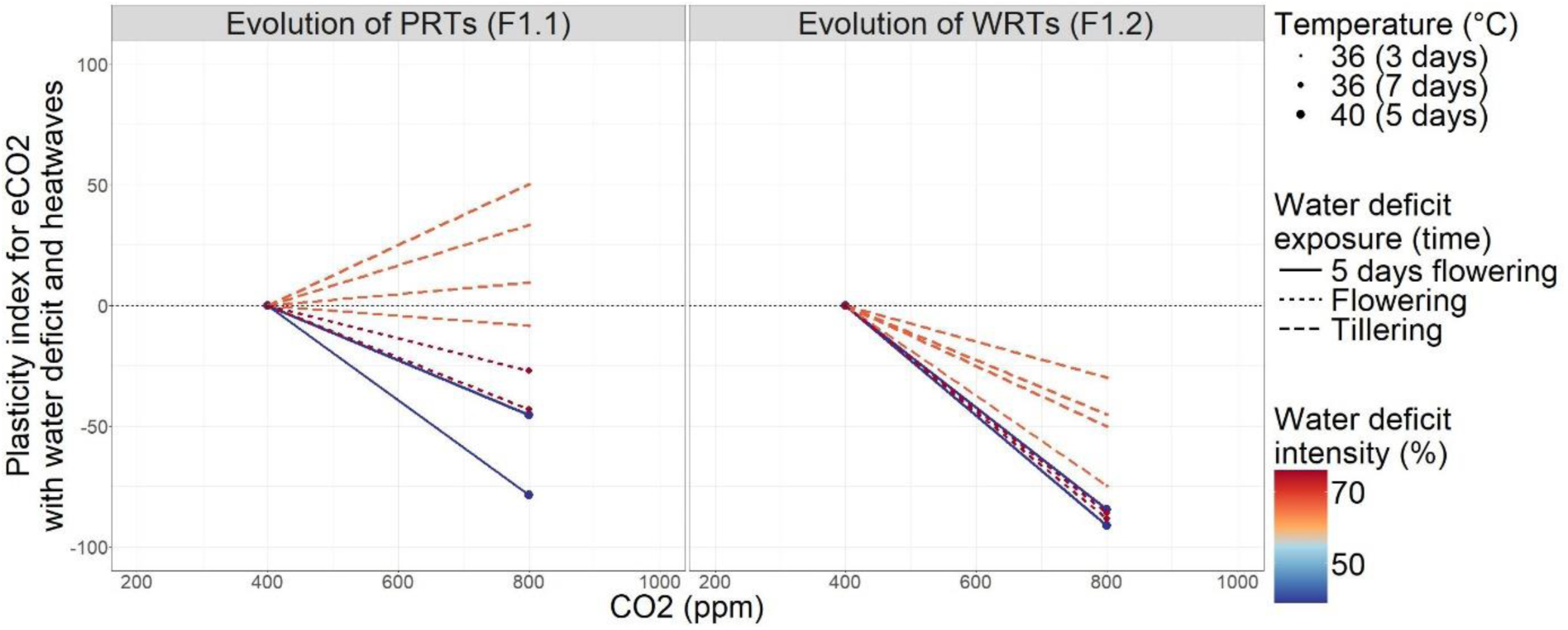
Relative reaction norms for the evolution of productivity (F1.1, Δ_[aCO2_×_WW_× *aT*_] vs [eCO2_×_WD_×HW_]_ (PRT), Eq 1) and water-related traits (F1.2, Δ_[aCO2_×_WW_× *aT*_] vs [eCO2_×_WD_×HW_]_ (WRT), Eq 1) for eCO_2_, water deficit and heatwaves. Productivity-related traits include photosynthesis, biomass and yield variables, water-related traits only include stomatal conductance variables. Results are presented for chamber facilities. Point sizes represent different heatwave temperatures, line types represent the onset of stress and colors represent different water deficit intensities (ratio of water inputs between treatments, in %).

With this approach we tackle the question of the compensation between eCO2 and climatic stress on productivity and water-related traits with focus F1 (brown arrows in Figure 2). We also tackle the question of interaction between eCO_2_ and stress on productivity-related traits by comparing eCO_2_ effect in stressed versus non-stressed conditions (purple arrows in Figure 2). In the rest of the review, the relative changes in plasticity indices are represented as relative reaction norms (Guntrip and Sibly, 1998) for the extracted observations, i.e. comparing the relative impact of a treatment (in %) with the control. No statistical analysis was performed on the plasticity indices due to the small number of observations and the heterogeneity of the data collected for crop productivity and water-related traits (e.g. studies performed at different scales from organ-to crop-level and from seconds to the whole crop cycle).

## 2. Combined effects of CO_2_ and water deficit

Water deficit and eCO_2_ both decrease stomatal conductance and transpiration (Figure 1), leading to improved water-use efficiency. However, they also have opposite effects on photosynthesis, which is decreased under water deficit but increased with eCO_2_ (Figure 1). Hence, plants display opposing plastic responses on photosynthesis under eCO_2_ and water deficit so plant biomass may increase or decrease depending on the characteristics of the water deficit and the level of CO_2_ increase. Zahra et al. (2023) concluded that eCO_2_ often compensates for the adverse effects of water deficit on grain development but the effects on grain quality are often detrimental with decreased grain size, protein content and grain protein yield. On another hand, both environmental factors trigger a plastic decrease in stomatal conductance, which is expected to decrease under combined eCO_2_ and water deficit, and this decrease might even be enhanced by their interactions. Such interactions can be studied for different concentrations of ambient and elevated CO_2_ as well as different intensities of soil water deficit estimated here as the percentage of water not applied in the water deficit treatment relative to the well-watered control (WI_WW vs WD_):

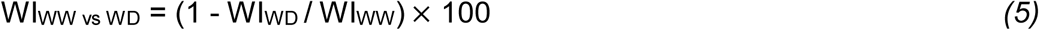

where WI is the water input (i.e. rainfall, irrigation, soil available water) under well-watered conditions (WW) or water deficit (WD) expressed as a percentage. Note that this water intensity index does not consider many characteristics that would influence stress intensity such as soil properties, soil available water at sowing, the level of evaporative demand, or the status of the plant when water stopped being applied to plants (as plants with greater leaf area typically deplete soil water faster than smaller plants).

### 2.1. Meta-information of the experimental studies

We identified 26 studies investigating the effect of interactions between eCO_2_ and soil water deficit on wheat (detailed in Table S1). The characteristics of these experiments are summarized in Table 1 for the studied ranges of CO_2_ concentrations, water deficit intensity, timing of stress, type of observations and number of cultivars.

**Table 1.** Characteristics of the 26 experiments analyzed that studied elevated CO_2_ x soil water deficit interactions in growth chambers, open-top chambers (OTC) and free-air CO_2_ enrichment (FACE), with the studied range of CO_2_ concentrations water deficit intensity (based on relative water input), the distribution of observations according to water deficit timing (i.e. entire growth cycle (continuous), pre-flowering (from 1^st^ leaf to flowering) or around flowering), and the distribution of observations for productivity and water-related traits. Productivity-related traits (PRTs) include yield components (grain yield, ear weight, ear dry matter or grain test weight per m^2^, per plant, pot or shoot and either at harvest or at the end of grain filling) or biomass/plant weight variables (total dry weight, shoot dry weight per plant). Water-related traits (WRTs) include stomatal conductance (at flowering or at day 226), transpiration (cumulative transpiration, water use) and WUE (for shoots or the whole canopy, i.e. both shoots and roots).

| Facility | Chamber | OTC | FACE |
| --- | --- | --- | --- |
| CO<sub>2</sub> range (ppm) | 200-1000 | 350-900 | 370-700 |
| Water deficit intensity range (%) | 19-100 | 25-50 | 44-58 |
| Number of observations | 63 (16 studies) | 9 (6 studies) | 7 (4 studies) |
| Timing of water deficit (starting point) | 16 continuous, 1 flowering, 46 pre-flowering | 5 continuous, 4 pre-flowering | 3 continuous, 2 pre-flowering, 2 flowering |
| PRTs | 63 (16 biomass, 47 yield) | 9 (2 biomass, 7 yield) | 7 (yield) |
| WRTs | 52 (42 stomatal conductance, 4 transpiration, 6 WUE) | 4 (2 stomatal conductance, 2 WUE) | 1 (stomatal conductance) |
| Number of cultivars | 30 | 5 | 6 |

### 2.2. Compensation between ambient and future-like climatic conditions combining eCO2 and water deficit (F1)

#### 2.2.1. Compensation for productivity-related traits (F1.1)

For the studied data, the difference in productivity-related traits (PRTs) between non-stressed conditions under aCO_2_ and stressful conditions under ‘future-like’ eCO_2_ (F1.1, Δ_[aCO2×WW] vs [eCO2×WD]_ (PRT), Eq 1) is negative in 70% (i.e. 55/79) of the considered observations. Reaction norms are displayed on Figure 3 (F1.1) gathering different types of traits (yield and plant biomass) and the detail per trait is available in Figure S1A. The relative difference has an average of −9% and a large variability (SD 33%; Figure 3 F1.1). There are more cases of short and/or severe water deficits (Figure 3, red-dashed lines) in chamber studies than in OTC or FACE studies and they tend to result in more negative impacts on productivity. For water deficit applied during part of the plant cycle only, stress onset was mostly applied before flowering, hence with longer exposure than for the few observations with an onset at flowering (3 green diamonds on Figure 3 F1.2). These three observations (Asif et al., 2017; Erice et al., 2014) corresponded to limited yield response (0 to −9%) for moderate stress intensities of 45-50%. In chambers, productivity response is more negative for punctual water deficits than continuous ones; this is likely due to higher water deficit intensities for these punctual deficits (80-100% versus 20-60% for continuous ones) and the fact that the duration of these punctual deficits is still quite long (start pre-flowering until end of growth). The existence of some positive reaction norms for low intensity water deficits shows that in such conditions eCO_2_ can compensate for water deficit and result in a positive productivity response. These positive responses are not systematically related to genetic characteristics: it is the case in some experiments using drought-tolerant cultivars as Mv15 in Harnos et al. (2002), Ganmai in Wu et al. (2004), Gladius in Cao et al. (2022). Some other genotypes also show positive productivity response as Yongling in Kang et al. (2002), Cham 1 in Kaddour and Fuller (2004), Soissons in Acker et al. (2024), Regallo in Medina et al. (2016) or Kolompos in Farkas et al. (2021); these cultivars are not described as drought-tolerant or exhibiting other traits that may suggest a tolerant behavior, they are often the most widely used in the regions where the experiments take place.

Overall, the impact of soil water deficit highly depends on the stress onset timing, duration and intensity, as well as other factors related to the soil and crop (Chenu, 2015), both under aCO_2_ (Tardieu et al., 2018) and eCO_2_ (Figure 3). Understanding the quantitative interactions between CO_2_ levels and water deficit is needed (F2) to model crop responses in projected future climate scenarios. In the studied experiments, eCO_2_ does not compensate for the effect of water deficit on crop productivity, but the large variability of the observations highlights the fact that this impact depends on stress onset timing, duration and intensity.

#### 2.2.2. Compensation for water-related traits (F1.2)

Reaction norms for water-related traits (WRTs) between ‘current’ non-stressed and ‘future-like’ stressed conditions (F1.2, Δ_[aCO2_×_WW] vs [eCO2_×_WD]_ (WRT), Eq 1) are displayed on Figure 3 (F1.2) gathering different types of traits (stomatal conductance, transpiration and WUE) and the detail per trait is available in Figure S1B. The trend is negative with 43/49 negative relative reaction norms for stomatal conductance and transpiration with an average of −27% (SD 21%), and only positive relative reaction norms for WUE with an average of +61% (SD 14%; Figure 3 F1.2, purple diamonds). However, even if the hypothesis of a decrease in water consumption in response to combined eCO_2_ and water deficit was mostly confirmed, several exceptions were also reported. Some positive relative reaction norms were surprisingly obtained in the studies by Medina et al. (2016) and Farkas et al. (2021), due to the increase in conductance in response to eCO_2_ exceeding the usual decrease due to water deficit, but these findings were not significant. Overall, the global trend is towards reduced water consumption under combined eCO_2_ and water deficit.

### 2.3. CO2 interaction with water deficit on productivity-related traits (F2)

The hypothesis of whether eCO_2_ has a stronger effect on productivity-related traits (PRTs) under dry than wet conditions (F2, Δ_eCO2, [WW] vs [WD]_ (PRT), Eq 2) has been tested by several studies with contrasted results. Reaction norms are displayed on Figure 3 (F2) gathering different types of traits (yield and plant biomass) and the detail per trait is available in Figure S1C. In the studied dataset, 54% (i.e. 43/79) of the relative reaction norms were positive, indicating a stronger effect of eCO_2_ under dry conditions in more than half of the studied combinations of water deficit and eCO_2_ (Figure 3 F2). However, the mean difference in this relative effect of eCO_2_ between well-watered and dry conditions is close to zero (1.83%) with a large variability indicated by a standard deviation of 44%. In other words, no clear trend could be observed due to this high standard deviation relative to the mean difference. Thus, the hypothesis that the positive effects of eCO_2_ are stronger under dry than under wet conditions was far from systematically confirmed for wheat.

## 3. Interactive effects between eCO_2_ and warming

Increased temperature decreases the specificity of Rubisco for CO_2_, favoring photorespiration with oxygenation rather than photosynthesis with carboxylation. Hence, the benefit of eCO_2_ is expected to be even greater under high temperatures due to heat inhibition of photosynthesis (Long, 1991; Scafaro et al., 2023). Photosynthesis models predict that for C3 plants, response to eCO_2_ might raise the optimum temperature of CO_2_ assimilation by +5°C, resulting in a positive interaction between these two variables (Alonso et al., 2009; Drake et al., 1997; Long, 1991) but this theory was not verified by all investigators (Qaderi and Reid, 2009). Nevertheless, when temperatures exceed the optimum, notably during heatwaves, photosynthetic activity decreases, counteracting the positive effect of high CO_2_ levels (Abdelhakim et al., 2022). Increases in CO_2_ and mean temperature also have opposite effects on transpiration, which decreases with eCO_2_ but increases with warming, to cool the leaves (Rivero et al., 2022). The contrasting responses to eCO_2_ and warming observed for photosynthesis and stomatal conductance are making it difficult to predict the net result of these effects. Kadam et al. (2014) found that reproductive physiology was driven mainly by temperature, with only an indirect impact of eCO_2_, leading to reduced grain set and yield. In addition, eCO_2_ may result in a slightly smaller increase in grain protein levels in heat-treated plants. Overall grain quality seems to decrease with increasing CO_2_ levels and warming, mainly due to changes in storage proteins under both these sets of conditions, decreasing dough and baking quality (Farooq et al., 2011). However, Zahra et al. (2023) concluded that eCO2 combined with warming would increase grain size as well as globulin and albumin content, resulting in increased grain protein content despite a decrease in gliadin and glutenin.

### 3.1. Meta-information of the experimental studies

We identified 32 studies focusing on interactions between eCO_2_ and warming, 23 of them studying warming through high temperatures (detailed in Table S2) and 9 studying warming through heatwaves (detailed in Table S3). The characteristics of these experiments are summarized in Table 2 for the studied ranges of CO_2_ concentrations, temperature increment or stress temperature, timing and duration of stress, type of observations and number of cultivars.

**Table 2.**
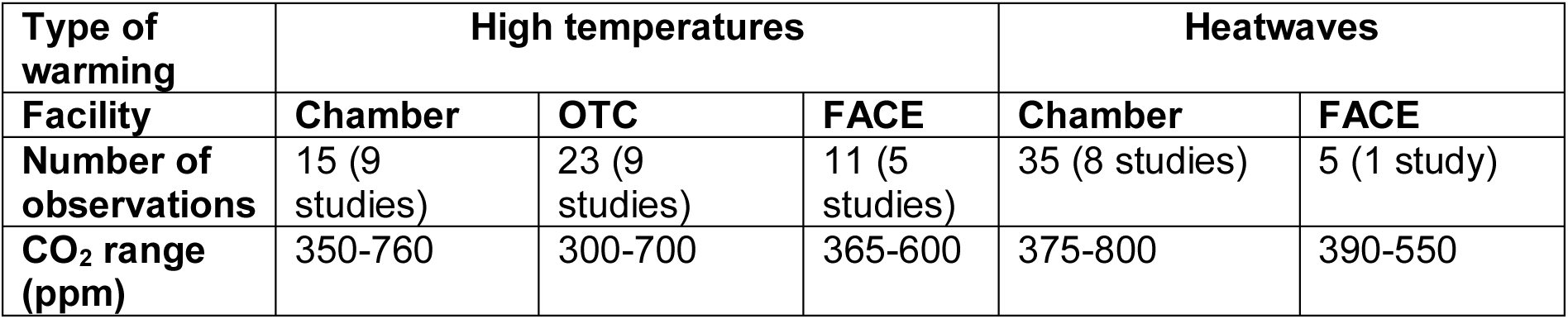
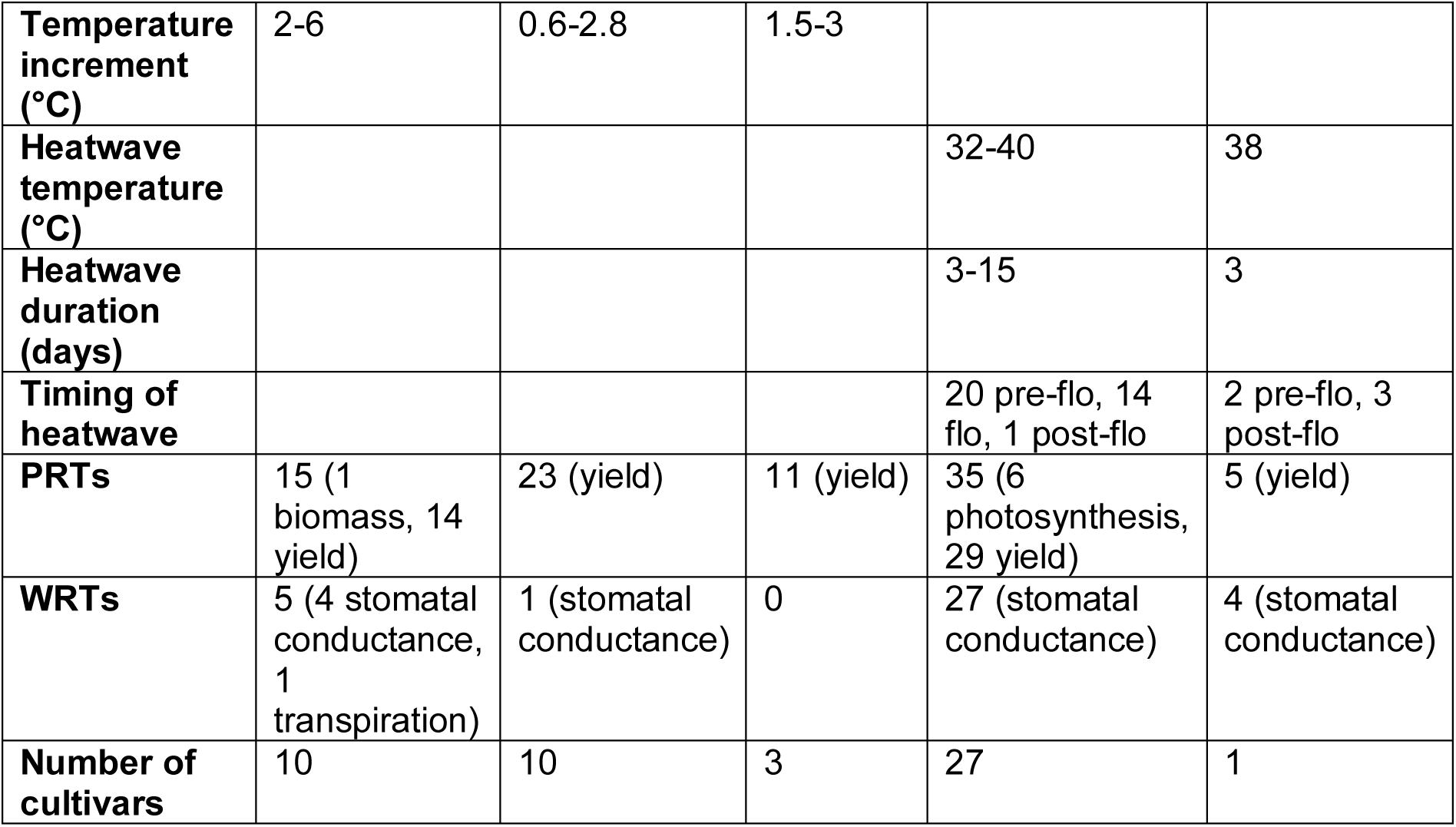
Characteristics of 32 experiments analyzed that studied elevated CO_2_ with warming through high temperatures or heatwaves (HW), reported by types of experiments, i.e. growth chambers, open-top chambers (OTC) and free-air CO_2_ enrichment (FACE), with the studied range of CO_2_ concentrations and the constant temperature increase throughout the growth cycle or throughout the grain-filling period (2 observations) for HT, or the heatwave mean temperature and stress duration, the onset of heatwave before flowering (from tillering to heading), during or after (from 10 days after flowering to late grain filling), and the distribution of observations for productivity- and water-related traits. Productivity-related traits include variables for yield (grain yield per m^2^, per ear, per plant or per plot for high temperatures and grain yield per plant, per plot or for 10 spikes for heatwaves), plant above-ground biomass (dry biomass at week 5 for high temperatures), and photosynthesis (photosynthesis at flowering for heatwaves). Water-related traits include stomatal conductance (stomatal conductance, stomatal conductance at flowering, stomatal conductance during grain filling for high temperatures and stomatal conductance for heatwaves) and transpiration (evapotranspiration for high temperatures).

| Type of warming | High temperatures |  |  | Heatwaves |  |
| --- | --- | --- | --- | --- | --- |
| Facility | Chamber | OTC | FACE | Chamber | FACE |
| Number of observations | 15 (9 studies) | 23 (9 studies) | 11 (5 studies) | 35 (8 studies) | 5 (1 study) |
| CO<sub>2</sub> range (ppm) | 350-760 | 300-700 | 365-600 | 375-800 | 390-550 |
| **Temperature increment (°C)** | 2-6 | 0.6-2.8 | 1.5-3 |  |  |
| **Heatwave temperature (°C)** |  |  |  | 32-40 | 38 |
| **Heatwave duration (days)** |  |  |  | 3-15 | 3 |
| **Timing of heatwave** |  |  |  | 20 pre-flo, 14 flo, 1 post-flo | 2 pre-flo, 3 post-flo |
| **PRTs** | 15 (1 biomass, 14 yield) | 23 (yield) | 11 (yield) | 35 (6 photosynthesis, 29 yield) | 5 (yield) |
| **WRTs** | 5 (4 stomatal conductance, 1 transpiration) | 1 (stomatal conductance) | 0 | 27 (stomatal conductance) | 4 (stomatal conductance) |
| **Number of cultivars** | 10 | 10 | 3 | 27 | 1 |

### 3.2. Compensation between ambient and future-like climatic conditions with eCO2 and warming (F1)

#### 3.2.1. Compensation for productivity-related traits (F1.1)

Reaction norms studying productivity-related traits (PRTs) between non-stressed conditions under ambient CO_2_ and stressed conditions under ‘future-like’ eCO_2_ when considering high temperatures (F1.1, Δ_[aCO2×aT] vs [eCO2×HT]_ (PRT), Eq 1) are displayed on Figure 4 gathering different types of traits (yield and plant biomass) and the detail per trait is available in Figure S2A. The trend is nearly null (+0.03%) and variable (SD 14%), with about as many positive (25/49) as negative (24/49) relative reaction norms (Figure 4). In chambers, productivity responses are the most variable, depending on the degree of temperature increment. Responses are negative for high increases of 4-6°C (Hansen et al., 2019; McKee and Woodward, 1994; Mitchell et al., 1995, 1993), indicating that eCO_2_ cannot compensate for the negative effect of temperature increase. However, there are also some negative response for medium temperature increments of +3°C (Asif et al., 2019; Hakala, 1998), even though mean high temperatures should not be stressful (18-20°C) but the results were contrasted over different years for cultivar Polkka in Hakala (1998). Among the tested cultivars, none was characterized for heat tolerance, leading to a part of variability in plant response. In OTCs and TGTs results were also variable but to a lesser extent due to the more limited range of temperature increase. Some cultivars were selected for their heat-tolerance and showed positive productivity response such as DBW14, HD2967 (Prakash et al., 2017) and Zhongmai 9 (Wang et al., 2018) but negative for Halna (Dwivedi et al., 2017). Some other genotypes showed clear negative responses (Minaret, Tanmai, Yangmai) or positive (Timai) but they were not characterized for heat-tolerance, and other cultivars showed variable responses depending on the year. Finally, for FACE studies, productivity responses are mostly positive due to low temperature increments, especially in Braunschweig, Germany (Krause et al., 2023; Manderscheid et al., 2020) and Walpeup, Australia (Fernando et al., 2012; Fitzgerald et al., 2016). However, the study conducted in Kangbo, China (Cai et al., 2016) for cv Yangmai resulted in negative productivity responses over two years, which could be explained by the heat-sensitivity of this cultivar.

When considering heatwaves (F1.1, Δ_[aCO2×aT] vs [eCO2×HW]_ (PRT), Eq 1), reaction norms are displayed on Figure 5 (F1.1) gathering different types of traits (yield and photosynthesis) and the detail per trait is available in Figure S2B. The relative difference between ambient and future-like conditions was more negative on average (−8.5%), but with about half of negative responses (21/40) and an important variability (SD 41%, Figure 5 F1.1). So, in the studied experiments combining eCO2 and heatwaves there were as many positive as negative combined effects. The FACE study of Macabuhay et al. (2018) reported consistent positive responses, for a short heatwave duration (3 days) with high temperatures (38°C) applied before or after flowering. Productivity was particularly enhanced in the second year by combined eCO2 and heat for low-yielding conditions (high temperatures, low water availability) for cv Yitpi; this cultivar showed positive responses in terms of productivity despite not being heat tolerant in Chavan et al. (2022). The temperatures studied in chambers were higher than those studied in FACE studies, and tended to generate more negative productivity trends. For experiments in chambers under eCO_2_ and heatwaves, heat stress before flowering (solid lines in Figure 5) led to more yield loss, even with short duration and under lower temperature (35°C) than some experiments with higher temperature but later stress onset (red relative reaction norms at 40°C). Hence in the studied experiments combining eCO_2_ and heatwaves, productivity tends to decrease, with a greater impact for heatwaves before flowering. Indeed, pre-flowering heatwaves can severely impact biomass production with reduced photosynthesis, even though the earlier the stress, the more likely compensations could occur later on. However early heat stress could be more harmful as younger plants might have lower defense. This confirms the findings of Ullah et al. (2024) that also concluded in stronger yield reduction under pre-flowering than post-flowering heatwaves, mostly due to the reduction in grain number. Nevertheless the effect of timing is likely interacting with genetic effects; for example the 3 days heatwave at 35°C at meiosis experimented in Bokshi et al. (2021) issued a more negative response than the 3 days heatwave at 40°C at flowering experimented in Ulfat et al. (2021), which could be explained by the difference in timing but also by the genotypes selected for the experiments as there are many heat-sensitive cultivars (PBIC, Vixen, Devil, CMSA, Cobra, Dart, Suntop, Scout, E Rock) in Bokshi et al. (2021) compared to Ulfat et al. (2021) with mostly tolerant cultivars (Chakwal-50, Shahkar, Pakistan-13, FSD-08). However, some positive response can be observed for sensitive cultivars and inversely.

#### 3.2.2. Compensation for water-related traits (F1.2)

For the studied data, the difference in water-related traits (WRTs) between non-stressed conditions under aCO_2_ and stressed conditions under ‘future-like’ eCO_2_ when considering high temperatures (F1.2, Δ_[aCO2×aT] vs [eCO2×HT]_ (WRT), Eq 1) exhibits considerable variability (SD 83%) due to the low number of observations (only six from one study including five observations on stomatal conductance and one on transpiration; Figure S3). Although the trend is slightly positive (+3%), it should be noted that out of these six observations, four were negative, one at 0% and one at 150% under combined eCO_2_ and high temperatures (Yang et al., 2023). In this study, plant water status was studied by measuring stomatal conductance at leaf scale, which tended to decrease in response to high temperatures in 2018 and 2019, with a large additional decrease in response to eCO_2_. However, in 2020, stomatal conductance increased considerably in response to eCO_2_ (+79% under ambient temperature and +32% under high temperature) but this surprising response was no adressed in the study.

When considering heatwaves, (F1.2, Δ_[aCO2×aT] vs [eCO2×HW]_ (WRT), Eq 2), reaction norms are displayed on Figure 5 (F1.2) for stomatal conductance. The difference in water-related traits between ambient and future-like conditions displayed a mean change of −10% (SD 56%), and most (20/31) relative reaction norms were negative (Figure 5 F1.2). There were four null relative reaction norms and seven positive ones, including four observations for which the positive relative reaction norms were explained an increase in stomatal conductance with heatwaves that exceeded its decrease caused by eCO_2_. The positive responses for the other three observations of conductance was due to an increase in response to both eCO_2_ and heatwaves. Two of the 13 observations reported by Bokshi et al. (2021) involved large changes of +85 and +230% due to major increases in stomatal conductance in response to eCO_2_ not highlighted in their study. These positive reactive norms do not necessarily reflect an effect of scale (leaf vs plant) as responses in absolute values were minor with low values for the control treatment (ambient CO_2_ and temperature). Stomatal conductance in expected to increase under high temperatures (Poudyal et al., 2019) as a mechanism to cool leaf temperature, opposing the effect of eCO_2_ that decreases conductance; the reviewed studies show an overall downward trend in stomatal conductance, irrespective of the timing and duration of heatwaves. However, the response of stomatal conductance to heat is highly variable, notably depending on the timing of the measure (during or after stress). As expected, some studies found that conductance increases under heat (Bokshi et al., 2021; Shanmugam et al., 2013) but others concluded in a decrease (Shokat et al., 2021; Zhang et al., 2018) while the others concluded in non-significant effects of heat (Chavan et al., 2022, 2019; Macabuhay et al., 2018).

### 3.3. CO_2_ interaction with warming on productivity-related traits (F2)

We also studied if the fertilizing effect of CO_2_ is stronger under warming (through high temperatures or heatwaves) than at ambient temperature (aT; F2; Figure S4). For high temperatures, Δ_eCO2_, _[aT] vs [HT]_ (PRT) displayed a mean value of 5.8% and was highly variable (SD 28%), with 69% (24/35) positive relative reaction norms corresponding to a stronger effect of eCO_2_ under high temperatures (Figure S4A). For heatwaves, Δ_eCO2, [aT] vs [HW]_ (PRT) displayed a mean value of 30% with considerable variability (SD 97%) with 55% positive relative reaction norms (22/40), corresponding to a stronger eCO_2_ effect under heatwaves (Figure S4B). These positive means and higher proportion of positive relative reaction norms are relevant with the hypothesis of greater eCO_2_ effect under high temperature (Long, 1991; Qaderi and Reid, 2009) but it is not systematically verified and given the high degree of variability of the relative reaction norms, no firm conclusions can be drawn concerning the effect of interactions between eCO_2_ and warming on plant productivity.

## 4. Interactive effects of CO_2_, warming and water deficit (three-way interaction)

The combined effects of warming (through high temperature or heatwaves) with water deficit appear to aggravate yield reductions compared to single-stress effects (Kadam et al., 2014). However, the combination of warming and drought stress with eCO_2_ has been rarely studied. In a general review of the effects of this combination, Abdelhakim et al. (2022) highlighted opposite impacts of each single factor on photosynthesis and transpiration (Figure 1). As mentioned above, photosynthetic activity is increased by eCO_2_ and can be increased under combined eCO_2_ and high temperatures over an optimal temperature range, but it is reduced by heat and drought stress. Stomatal conductance (and hence relative transpiration) decreases under eCO_2_ and water deficit, thus allowing water saving, but increases under heat stress to allow leaf cooling. These individual effects have been well studied (Figure 1) but competing responses to eCO_2_, water deficit and high temperatures observed for photosynthesis and stomatal conductance lead to a difficult prediction of these responses when these stresses are combined. Overall, more needs to be done to understand properly the interacting effects in response to eCO_2_, water deficit and high temperatures interactions with dedicated experiments and meta-analyses based on historical data. Helman and Bonfil (2022) found that the alleviation effect of eCO_2_ of the last six decades has been insufficient to compensate for the global negative effects of heat and drought stress on yield. However, with the rate of climate change, such studies cannot be used to predict the future and studies based on simulation outputs provide contradicting predictions (Hochman et al., 2017; Makowski et al., 2020).

### 4.1. Meta-information of experimental studies

We identified 5 studies focusing on interactions between eCO_2_, warming and water deficit; 2 of them studying warming through high temperatures (detailed in Table S4) and 3 studying warming through heatwaves (detailed in Table S5). The characteristics of these experiments are summarized in Table 3 for the studied ranges of CO_2_ concentrations, temperature increment or stress temperature, timing and duration of stress, type of observations, intensity and timing of water deficit and number of cultivars.

**Table 3.** Characteristics of 31 experiments analyzed that studied elevated CO_2_ with warming through high temperatures or heatwaves (HW), reported by types of experiments, i.e. growth chambers, open-top chambers (OTC) and free-air CO_2_ enrichment (FACE), with the studied range of CO_2_ concentrations and the constant temperature increase throughout the growth cycle or throughout the grain-filling period (2 observations) for HT, or the heatwave mean temperature and stress duration, the onset of heatwave before flowering (from tillering to heading), during or after (from 10 days after flowering to late grain filling), and the distribution of observations for productivity- and water-related traits. Productivity-related traits include variables for yield (grain yield per m^2^, per ear, per plant or per plot for high temperatures and grain yield per plant, per plot or for 10 spikes for heatwaves), plant above-ground biomass (dry biomass at week 5 for high temperatures), and photosynthesis (photosynthesis at flowering for heatwaves). Water-related traits include stomatal conductance (stomatal conductance, stomatal conductance at flowering, stomatal conductance during grain filling for high temperatures and stomatal conductance for heatwaves) and transpiration (evapotranspiration for high temperatures).

| Type of warming | High temperatures |  | Heatwaves |
| --- | --- | --- | --- |
| Facility | OTC/TGT | FACE | Chamber |
| Number of observations | 8 (2 studies) | 13 (3 studies) | 8 (3 studies) |
| CO<sub>2</sub> range (ppm) | 360-700 | 360-550 | 400-800 |
| Temperature increment (°C) | 2-6 | 0.5-5.8 |  |
| Heatwave temperature (°C) |  |  | 36-40 |
| Heatwave duration (days) |  |  | 3-7 |
| Timing of heatwave |  |  | 4 pre-flowering, 4 post-flowering |
| Water deficit intensity (%) | 100 | 23-100 | 39-76 |
| Water deficit timing | 8 at flowering | 13 continuous | 4 pre-flowering, 4 flowering |
| PRTs | 8 (yield) | 13 (yield) | 8 (4 photosynthesis, 2 biomass, 2 yield) |
| WRTs | 2 (transpiration) | 1 (transpiration) | 8 (stomatal conductance) |
| Number of cultivars | 4 | 3 | 8 |

### 4.2. Compensation for productivity-related traits between ambient and future-like climatic conditions with combined eCO2, water deficit and warming (F1.1)

For the studied data, the difference in productivity-related traits between non-stressed condition under aCO_2_ and stressed conditions under ‘future-like’ eCO_2_ when considering water deficit and high temperatures (F1.1, Δ_[aCO2×WW×aT] vs [eCO2×WD×HT]_ (PRT), Eq 1) or heatwaves (Δ_[aCO2×WW×aT] vs [eCO2×WD×HW]_ (PRT)) is negative with mean values of −25% (SD 20%) for interactions with high temperatures (Figure 6 F1.1 for yield) and −14% (SD 43%) for interactions with heatwaves (Figure 7 F1.1 gathering yield and photosynthesis, details per trait in Figure S5). For interactions with high temperatures, most relative reaction norms were negative (18/21), likely due to the high intensity of water deficit with no irrigation and high temperature increments above +2°C in most studies. In FACE studies, results were less negative for the observations made in China (Guoju et al., 2005) than for the ones made in Australia (Fitzgerald et al., 2016; O’Leary et al., 2015) but temperature increments and water deficit intensity were also lower in China. Some studies investigated different temperature increments and found that the higher the increase in temperature, the lower productivity under combined eCO_2_, water deficit and high temperatures (Dias De Oliveira et al., 2013; Guoju et al., 2005). Dias De Oliveira et al. (2013) found that two cultivars with different vigor had similar productivity responses because cv Janz could compensate its lack of vigor by increased tillering capacity. Hence, Dias De Oliveira et al. (2015) investigated the role of tillering capacity and they found that high tillering capacity enabled enhanced productivity under combined eCO_2_ with water deficit and high temperatures, concluding that tillering is a promising target for breeding under adverse climatic conditions.

For heatwaves, 5/8 reactions norms were negative but some productivity responses were positive, showing an increase up to 50% for photosynthesis (Figure S5), for which the direct effect of eCO_2_ resulted in a stronger alleviating effect. This could result from both the lower temperature intensity and duration of the heatwave and the later occurrence of water deficit at flowering in Abdelhakim et al. (2021). Besides, eCO_2_ more directly compensates photosynthesis than biomass and yield for which it is only indirect, so eCO_2_ alleviation could be more important for photosynthesis. Under combined eCO_2_, warming and water deficit, productivity tends to decrease due to an insufficient alleviation effect of eCO2. For these studies, drought-tolerance enabled less negative responses for cv Gladius compared to Paragon in Li et al. (2019). On the contrary heat-tolerant cultivars GN5 in Abdelhakim et al. (2021) and LM20 in Abdelhakim et al. (2022) displayed negative responses whereas heat-sensitive LM19, KU10 and SF29 displayed positive responses. This entails that heat-tolerance might not be an interesting feature under combined eCO_2_ and stress, unlike drought-tolerance; this could be explained by the stronger effect of drought than heat stress on plants (Zandalinas et al., 2021).

### 4.3. Compensation for water-related traits between ambient and future-like climatic conditions with combined eCO2, water deficit and warming (F1.2)

The difference in stomatal conductance (the only WRTs studied) between non-stressed conditions under aCO_2_ and stressed conditions under ‘future-like’ eCO_2_ when considering water deficit with either high temperatures (F1.2, Δ_[aCO2×WW×aT] vs [eCO2×WD×HT]_ (WRT), Eq 1) or heatwaves (Δ_[aCO2×WW×aT] vs [eCO2×WD×HW]_ (WRT)) was negative, with mean values of −23% (SD 9%) for combined eCO_2_, water deficit and high temperatures (Figure 6 F1.2) and −69% (SD 23%) for combined eCO_2_, water deficit and heatwaves (Figure 7 F1.2). This indicates that the conductance-lowering effects of eCO_2_ and drought were stronger than the increasing effect of warming. However, these conclusions are drawn from a very limited number of relative reaction norms.

## 5. Limitations and opportunities to study the effects of climate change in wheat crops

### 5.1. Contributions of experiments with interacting environmental factors

In this review, we used relative reaction norms to assess the plasticity of productivity- and water-related traits involved in the response to eCO_2_ combined with warming and/or water deficit. By evaluating experimental studies, we were able to outline major trends and identify knowledge gaps. The key findings of this review are summarized in Figure 8. The relative reaction norms for productivity- and water-related traits extracted from experimental studies revealed variable responses to eCO_2_, warming and water deficit, but several trends nevertheless emerged. We found that future-like conditions tested in experiments are expected to decrease productivity, with this decrease accentuated by greater water deficit intensity, temperature increase or duration of heatwave events. For the studied conditions, there is a strong overall synergistic negative effect of combined heat and water deficit on plant performance, which cannot be compensated by eCO_2_ (Figure 8A). These findings are in accordance with the conclusion of the meta-regression of Helman and Bonfil (2022) on historical data and with the meta-analysis conducted by Zhu et al. (2023). More specifically, from inferring eCO_2_ and high temperature response in OTC experiments, Zhu et al. (2023) found that eCO_2_ and high temperature projected in the future would result in global yield gain under a moderate emission scenario (Representative Concentration Pathway RCP 4.5) but in a global yield loss under a high emission scenario (RCP 8.5); by contrast, when inferring results from FACE data, eCO_2_ and high temperature projected by both RCP scenarios would always result in yield loss. We also found that heatwaves, despite their large diversity, had greater effects on productivity- and water-related traits than high temperatures when combined with eCO_2_. They should therefore be characterized with greater care in terms of intensity, timing and duration. In the considered studies, heat stress occurring before flowering seemed to be more harmful than for those applied later on but only in chambers for a temperature of 35°C, as there was not enough data in FACE to confirm this observation. However, such high temperatures rarely occur in wheat production environments (e.g. Ababaei and Chenu, 2020).

**Figure 8.**
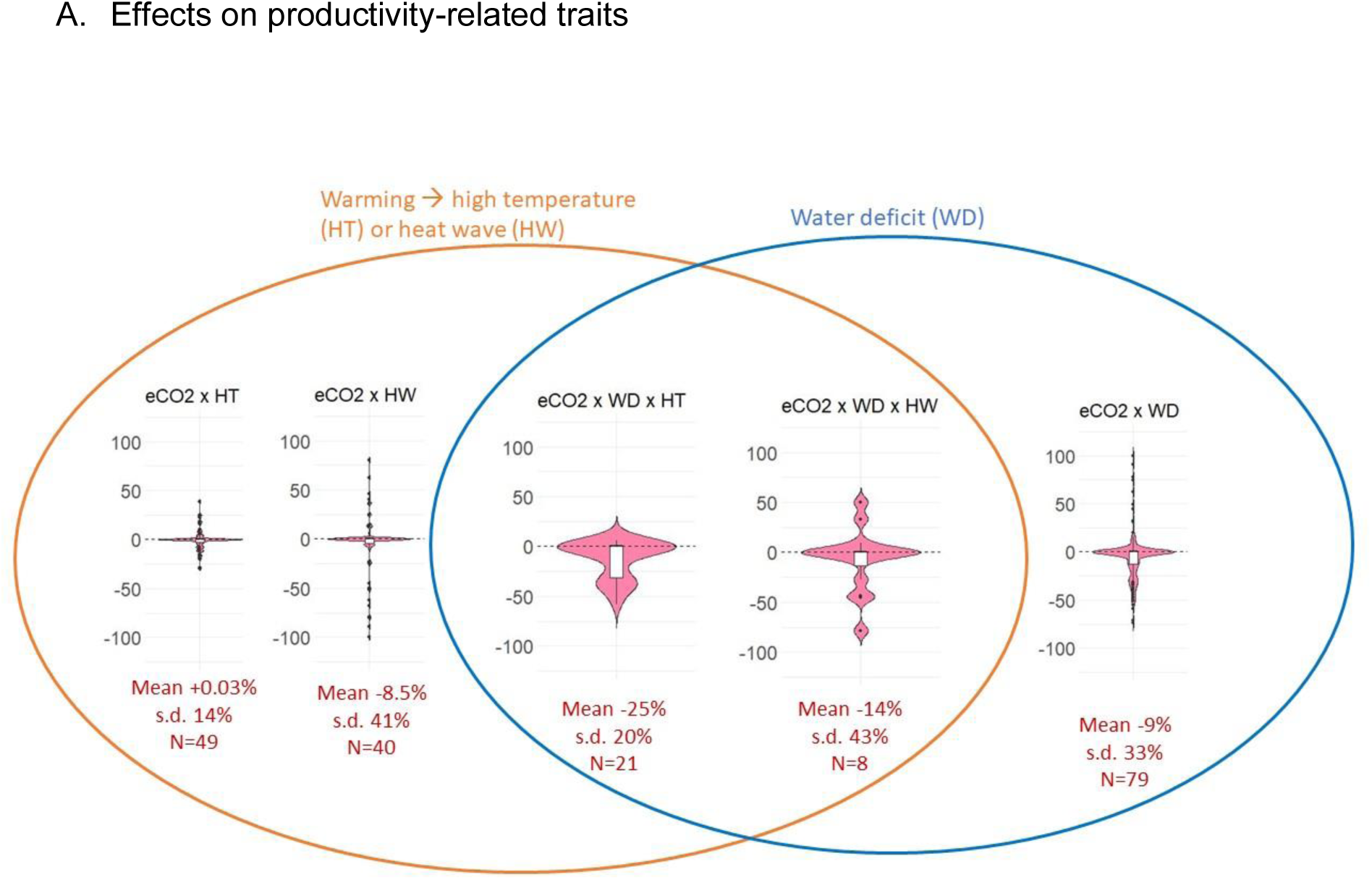
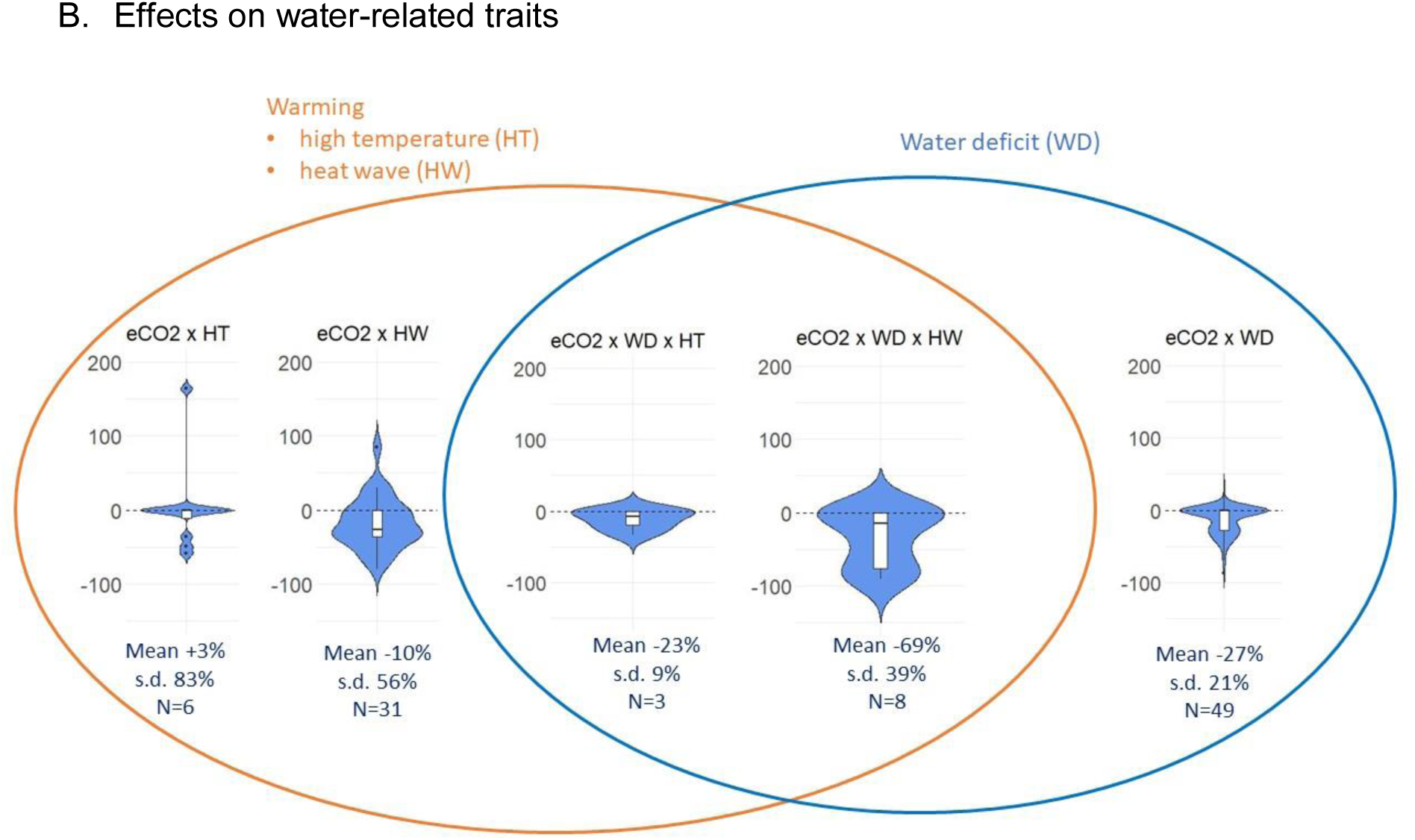
Summary diagram of plant plasticity for A) productivity-related traits (PRT, i.e. yield, plant biomass and photosynthesis in pink) and B) water-related traits (WRT, i.e. stomatal conductance and transpiration in blue) in response to elevated CO_2_ (eCO_2_), warming though high temperature (HT) or heatwave (HW), and water deficit (WD) for published experimental data (i.e. mix of stress with a wide range of intensity, timing and duration). Violin and box plots represent the distribution of the plasticity indices PI_[aCO2 x nS] vs [eCO2 x S]_, i.e. the relative variation in studied traits between non-stressed conditions under ambient CO_2_ (aCO_2_ x nS) and stressed conditions under ‘future-like’ eCO_2_ (eCO_2_ x S). The mean value, the standard deviation and the number of observations (N) are displayed below each graph.

It is often hypothesized that eCO_2_ has a greater effect in dry than in wet conditions (Wang et al., 2022), but only 54% of the 79 relative reaction norms calculated here for experiments focusing on both eCO_2_ and water deficit confirmed this hypothesis (Figure 3 F2), which is far from being systematically verified. The contrasting conclusions of these studies highlight the challenge to study quantitative response to combined stress in conditions that vary for those stresses (timing, intensity, duration of the stresses) and other factors (e.g. radiation, temperature, fertilization, genotypes). Stressful conditions under eCO_2_ could be expected to result in a global decrease in water consumption (e.g. Figures 3 F1.2, 5 F1.2, 6 F1.2 and 7 F1.2), even with the antagonistic effects of higher temperatures. This implies that the plastic increase in water consumption in response to warming is largely surpassed by the plastic decrease in response to eCO_2_ and water deficit. This finding is consistent with that of Zandalinas et al. (2021) concerning the stronger effect of drought than of heat stress on stomatal conductance responses. However, there were several surprising observations of increases in stomatal conductance in response to eCO_2_. Most of them were not significant but questioned a potential scale effect. Indeed, a scale effect (organ vs plant level) on the impact of eCO_2_ on stomatal conductance/transpiration has already been reported, with stomatal conductance decreasing in leaf tissues while transpiration of the whole plant increased due to the fertilization effect of eCO_2_ on biomass allowing plants to produce more leaf area. For instance in ryegrass, Acker et al. (2023) reported a decrease in stomatal conductance in leaf tissues in response to eCO_2_ but an increase in transpiration at plant scale. The studies reviewed here included observations at organ and plant scales, with a decrease in stomatal conductance/transpiration in response to eCO_2_ observed at both scales.

For a same combination of stress factors, relative reaction norms were found positive or negative depending on the intensity/timing/duration of the stress, highlighting the complexity of the interactions involved, and the fact that the combined effects of environmental variables may not be linear. More complex relative reaction norms can be drawn for studies considering more levels and modalities of environmental factors. For instance, two studies (Farkas et al., 2021; Varga et al., 2017) considered different levels of CO_2_ concentrations (400, 700 and 1000 ppm), water deficit (water input reduced by 100 and 82%, respectively) and stress onsets for five cultivars presented in Figure S6. These studies clearly highlight that yield and transpiration response to combined eCO_2_ and water deficit is not linear. This non-linearity notably questions the use of linear relative reaction norms to represent the effects of such stress. It suggests more complex frameworks should be developed, such as random regression mixed models enabling to analyze non-linear response (Arnold et al., 2019) or eco-physiological response curves of key traits derived from appropriate levels of stress, which can be integrated to crop models in order to account for interactions within the crop and with other environmental variables (Chenu et al., 2017; Collins et al., 2021; Hammer et al., 2010).

### 5.2. Limitations concerning the studied experimental set-ups

The experimental set-ups used were subject to limitations even for studies of the effects of eCO_2_, warming or water deficit separately. Enclosure studies tend to overestimate the effects of eCO_2_ due to their small scale and warmer temperatures (Long et al., 2004), whereas FACE studies, which assess eCO_2_ effects in the field, suggest a smaller effect of eCO_2_ on yield (Van Der Kooi et al., 2016). However, the large fluctuations of CO_2_ occurring within the day in FACE studies (due to intermittent CO_2_ emission and wind) may lead to an underestimation of the effects of eCO_2_ (Allen et al., 2020). Another downside of FACE studies is their cost. There is, therefore, currently no perfect experimental set-up for such studies. As a result, the CO_2_ concentrations tested rarely exceed 700 ppm, whereas projected CO_2_ concentrations range from 540 to 1300 ppm (Calvin et al., 2023).

Some environmental stresses are associated as they often occur together (e.g. heatwaves typically happen during drought periods) and have intertwined effects. For instance, the decrease in transpiration in response to a warming treatment can be due to the increase in atmospheric water deficit triggered by higher temperatures (Lobell et al., 2013; Sadok et al., 2021). Atmospheric drying should receive more attention in studies of water deficit (Wright and Collins, 2024) and warming.

We identified several limitations of the studied experiments combining eCO_2_, water deficit, and warming (through high temperatures or heatwaves) in this review. Firstly, water deficit was mostly studied over long periods, whether continuously from sowing to harvest or from sowing to flowering, the only exception being a four-day water deficit after flowering in one study (Li et al., 2019). These long water deficits include drought stress after flowering but are not representative of short occasional drought stresses that are frequent in production environments, particularly after flowering (Chenu et al., 2013). There is a gap in our knowledge concerning terminal drought during grain filling, due to a lack of studies on finite periods of drought stress at different times and for different durations, despite the major threat to crops posed by terminal drought (Farooq et al., 2014). Furthermore, these studies only considered soil water deficit, despite the importance of effects from atmospheric water deficit than often co-occur with soil water deficit and warming. Soil water deficit is also hard to compare between studies, as water deficit does not only depend on water input (as characterized here) but also on soil properties (e.g. soil water content at sowing, capacity of the soil to retain water), evaporative demand (driver of soil water loss) and plant characteristics (e.g. the feedback of plant growth on soil water depletion). For high temperatures and water deficit, we considered the increase in temperature or ratio of water input compared to control experiments, but temperature from the control and soil water availability are also important and were not always provided. A lot of studies were also conducted in relatively small pots, which can constrain root growth and typically have higher soil temperature (especially for warming studies) than in field conditions. Only very few studies consider different levels of stress (five for combined eCO_2_ with water deficit, one for eCO_2_ with high temperatures, two for eCO_2_ with water deficit and high temperatures and one for combined eCO_2_ with water deficit and heatwaves), while it is necessary to better understand the non-linearity of plant response to combined environmental factors. The range of CO_2_ concentrations studied was wider for studies focusing on water deficit (200-1000 ppm) than for those investigating warming (300-750 ppm), and was narrower in FACE studies, limiting long-term predictions. Nitrogen is an additional key factor that should be considered in the framework of climate change, notably due to the dilution effect of eCO_2_, so modifications in nitrogen fertilization practices in farming might be required. Finally, the focus of this review is on plant plasticity observed in experiments, but the relevance of the climatic perturbations generated in these experiments in regards to foreseeable climatic investigation should be more thoroughly investigated. Zhu et al., (2023) notably highlighted that the CO_2_ and mean temperatures ranges under future climatic projections are relevant with experimental variations in OTC facilities under RCP 4.5 but disjointed under RCP 8.5 late century, and more alarmingly the narrower ranges explored by FACE facilities are disjointed from climatic projections under both scenarios mid and late century. Hence experimental knowledge does not cover future possible climatic conditions, especially in FACE. It is hence critical that future experiments tackling combinations of eCO_2_, high temperature and water deficit focus on exploring larger climatic ranges relevant with climatic projections to gain better insight on their effect on wheat performance.

Relative reaction norms for productivity- and water-related traits between non-stress conditions with ambient CO_2_ and stressed conditions with future eCO_2_ levels were highly variable due to considerable data heterogeneity associated with differences in experimental protocols. We tried to take some of this variability into account, for the timing, intensity and duration of the stress and the type of variable considered (yield vs. plant biomass vs. photosynthesis and stomatal conductance vs. transpiration vs. WUE). However, there were very few observations by category, and yet this characterization remains coarse in resolution, only considering broad range of stress intensity/onset/duration within categories. Variability in the results also most likely arise from (i) other climatic factors that were not considered but affect plant growth and development, and from the fact that (ii) all these studies included a large number of genotypes rarely common between studies, some differing for heat or drought tolerance, or in specific traits (e.g. tillering capacity, vigor), with often a preference for local genotypes.

Transpiration and/or stomatal conductance should be more systematically measured, and more specific studies of the transpiration response to the combination of stresses under eCO_2_ need to be carried out. This will allow a better estimation of changes in canopy energy balance and consequently the increased plant organ temperature compared to air temperature. The more systematic acquisition of additional traits, such as tillering and leaf area, would improve our understanding of the effects of different stress factors.

There is currently no consensus for experimentally studying the effects of climate change on crops. Hence, the existing experimental data tackling combinations of eCO_2_, warming and water deficit is quite heterogeneous. This heterogeneity is interesting and informative because it represents to a certain extent a part of an important variability in real-life conditions across the globe. Thorough testing across a wide range of environments could appear as a solution but this heterogeneity will always be difficult to manage, and the number of conditions will never be as comprehensive as to give possible scenarios. We rather recommend considering the following key points by order of importance:

- The overall number of experiments combining eCO_2_, warming and water deficit should be increased, even if they remain heterogeneous, and experiments in small pots should be avoided.
- The studied ranges of climatic variations should be widened, especially for CO_2_ in FACE facilities even though the cost is high.
- More generally, experimented climatic variations should be better rooted in future climatic projections to reproduce likely future climatic patterns. Ecoclimatic indicators or stress index computed from crop models to account for interactions between plants and their environments might also allow to better characterize stress occurring in the experiments (Caubel et al., 2015; Chenu, 2015; Le Roux et al., 2024); they could be calculated for current and projected climate scenarios to design experiments relevant to the target population of environments (Chenu, 2015; Chenu et al., 2013b). More broadly, the quality of such experimental data could be enhanced by improving the design of the experiments, as recommended by Moshelion et al. (2024) for drought experiments for example.
- Even if experimental conditions remain heterogeneous, some common elements could be defined for experimental protocols, notably with more systematic and standardized measurements of traits underlying yield and water consumption. The use of common ecoclimatic indices for stress characterization could also be relevant for the comparison of these heterogeneous experiments.
- Another focal point for future studies should be the genetic response of plants to climate change, notably response to environmental factors such as CO_2_. Cultivars used in the experiments considered in this review were very diverse, some selected for their tolerance or sensitivity to heat and/or drought so they may differ in their plasticity to environmental variation, introducing more noise in the comparison of their relative reaction norms.

This will help in better understanding the possible compensations between CO_2_ and climatic stress, and the way they interact with each other. Improving the quality of this experimental data is also of major interest for crop modeling.

### 5.3. Perspectives for crop modeling

Crop models are efficient tools for overcoming the lack of data from studies of climate fluctuations and exploring the effects of these fluctuations on different processes. They are widely used for predictions under future climate scenarios (Chenu et al., 2017; Morel et al., 2021; Wang et al., 2023, 2017; Watson et al., 2017), their conclusions varying with the local context. Crop models are also useful for testing adaptation strategies, mainly based on the modulation of sowing date and genotype selection (Zheng et al., 2018; (Collins and Chenu, 2021; Zheng et al., 2018).

One key issue in the use of crop models to study climate change is their lack of validation with experimental data under combined fluctuations of CO_2_, temperature and water availability. Indeed, few validations have been reported to date. Ewert et al. (2002) compared crop simulations with FACE data obtained under combined eCO_2_ and water deficit conditions by Kimball et al. (1995), which gave realistic results. Asseng et al. (2019) studied an ensemble combining 32 wheat crop models and found that the model ensemble realistically simulated FACE data under combined eCO_2_, high temperatures and water deficit published in Fitzgerald et al. (2016), however the validity of individual models was not investigated. Considerable variability between wheat crop models has also been reported (Ahmed et al., 2019). Thus, even if the median of such model ensembles appears to be valid, some models may under- or overestimate key variables. Rezaei et al. (2023) imputed this uncertainty in model projections to differences in (i) model structure, (ii) the consideration of eCO_2_ effects, (iii) genotypic differences and (iv) methodologies for developing modeling routines to study climate change impact on crops. Crop models are rarely subjected to a quantitative validation against experimental data combining fluctuations of CO_2_, temperature and water availability, but empirical qualitative validation is more common. Better validations of crop models with field-based experimental data under climatic fluctuations are required. The lack of validation of crop models under such climatic conditions on existing data, is due partly to (i) limited availability of quality field data combining eCO_2_, water deficit and/or warming, as highlighted in this review, and (ii) the fact that FACE data explores a very limited range of climatic variations. Additional experimental data exploring wider climatic ranges is required for the use of crop models in future climatic projections. To do so, relevant dedicated methods should be considered. Some studies have concluded that model predictions are consistent up to CO_2_ concentrations of 500-700 ppm (Chenu et al., 2017; Toreti et al., 2020; Vanuytrecht and Thorburn, 2017). Other studies concluded that crop models overestimate CO_2_ effects (Addy et al., 2021; Ahmed et al., 2019), possibly due to model calibration on data from enclosure studies or a lack of consideration of acclimation to eCO_2_ in crop models, i.e. the inhibition of processes like photosynthesis under prolonged exposure to eCO_2_ but with little systematic evidence under field conditions (Tausz-Posch et al., 2020). Determining the validity of diverse crop models for different plant processes would facilitate the definition of appropriate modeling strategies and widen the range of validity of crop models under climate change. For longer-term predictions, crop models should also be calibrated on experimental data with higher CO_2_ concentrations (above 750 ppm), which is lacking for FACE studies (max CO_2_ level typically 600-750 ppm).

Genetic information related to adaptive traits should also be integrated to the simulated effects of CO_2_, water deficit, warming and other climatic factors in crop models through parametrization. This is an important issue of increasing interest as yield response to multiple stresses is cultivar-specific (Abdelhakim et al., 2021), although experimental data for calibration-validation of models for different genotypes are lacking.

In conclusion, better data quality is required for the validation of crop models in conditions combining eCO_2_, warming and water deficit. This will enable to strengthen and widen the range of validity of these models and make more robust predictions. Besides, confronting a model with experimental data is an essential practice that should be continuously performed to challenge the hypotheses implemented in the model, as the point is to learn the ways in which a model is false (Gelman and Shalizi, 2013). Indeed, understanding why a hypothesis implemented in a model is false enables to modify this hypothesis and formulate new ones and improve our general understanding of complex processes.

## 6. Conclusion

To better understand the interaction effects of eCO_2_, high temperatures, heatwaves and water deficit on crop functioning and performance, we reviewed the existing knowledge for these interactions and analyzed experimental data related to crop productivity and water consumption. Relative reaction norms for productivity- and water-related traits were highly variable due to the diversity of experimental protocols. Nevertheless, overall, productivity and water consumption in stressed conditions with elevated CO_2_ tended to be lower than in unstressed conditions with current ambient CO_2_. Also, the effect of eCO_2_ on productivity was not systematically stronger in drier conditions. Experimental studies on interaction between climatic variables are still lacking. Given the cost of such experiments, there is an urgent need to define common protocol elements to allow improved comparisons and development of meaningful databases, and this review suggests some guidelines to do it. Such experimental results are precious for calibration and validation of crop models, which are widely used to assess and adapt to impacts of future climate despite not having thoroughly tested for such applications.

## Acknowledgments

We thank R. Barillot, C. Furusho-Percot and D. Cammarano for providing feedback for some of the results obtained in this review, and J. Wolf for providing data. We also thank the FSOV REGARD project for funding this work.

## Supplementary Material

**Table S1.**
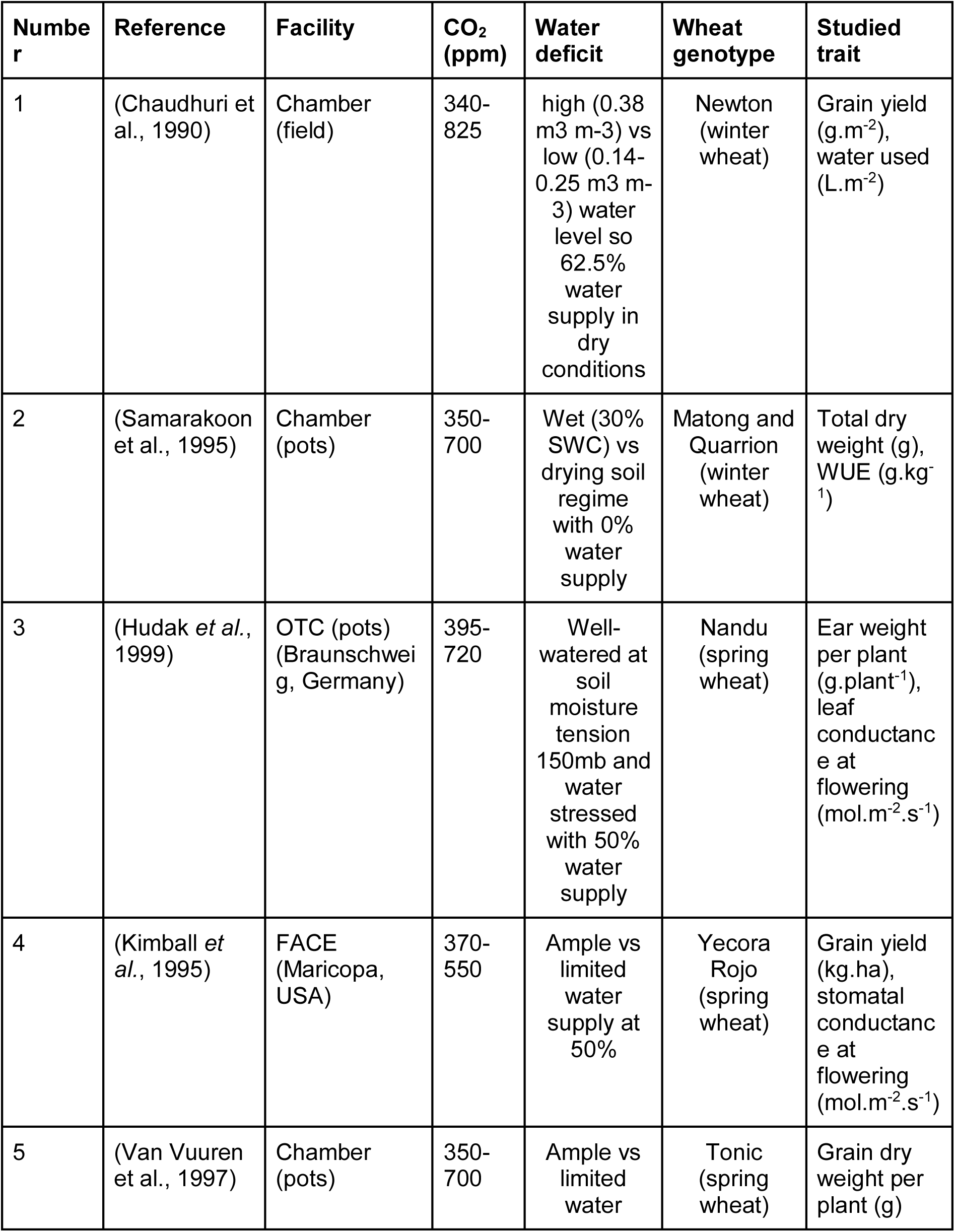
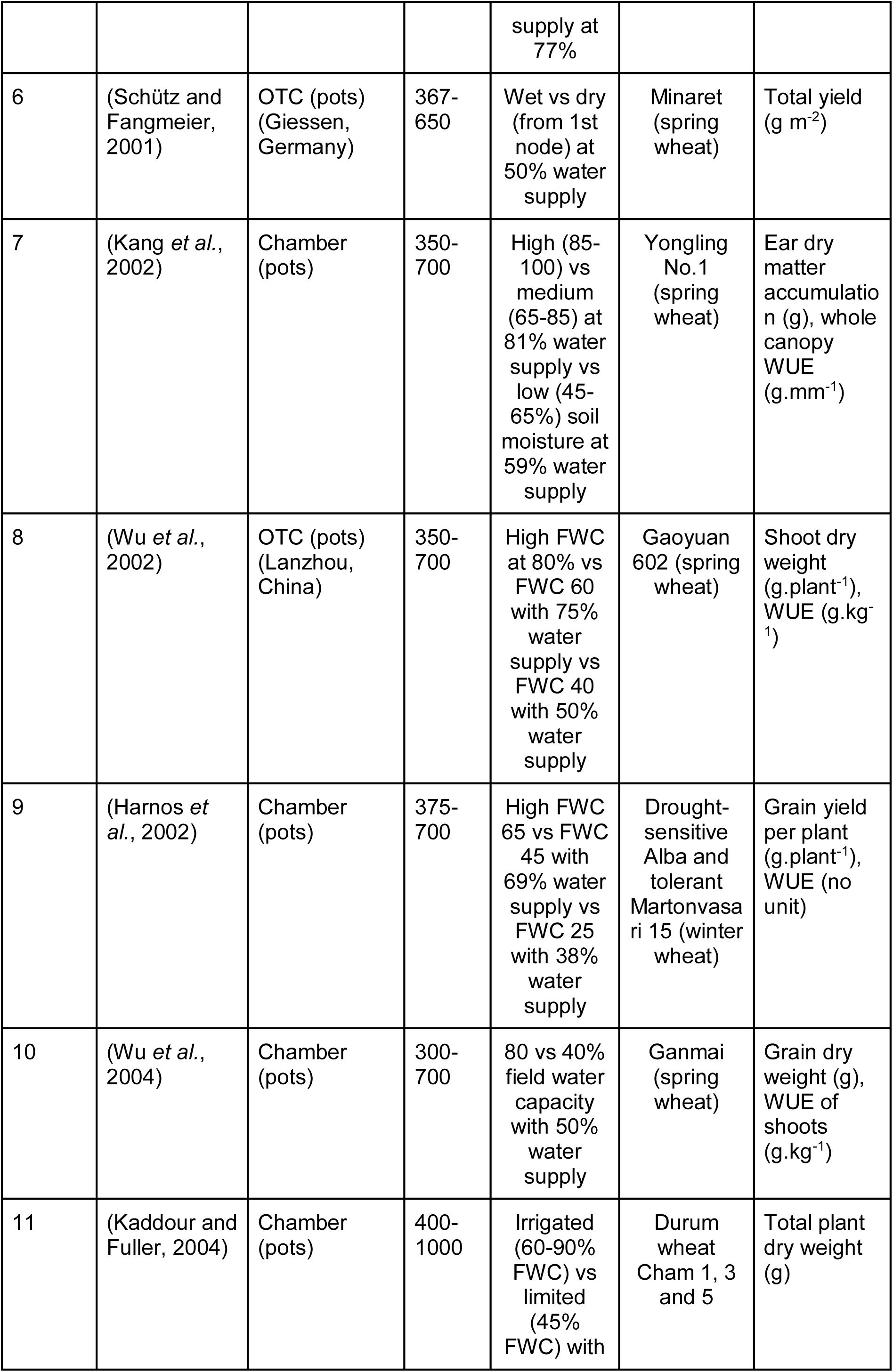
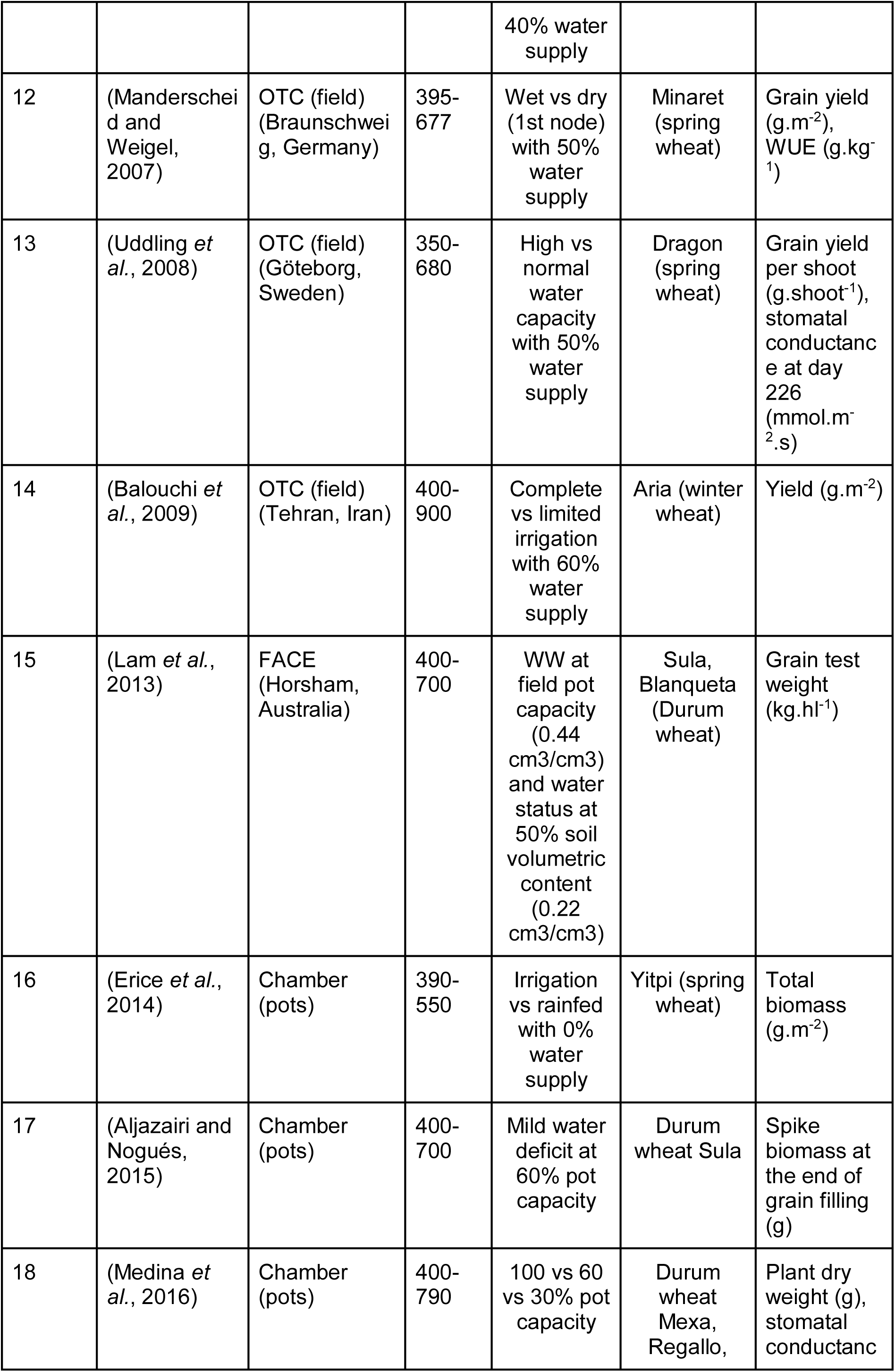
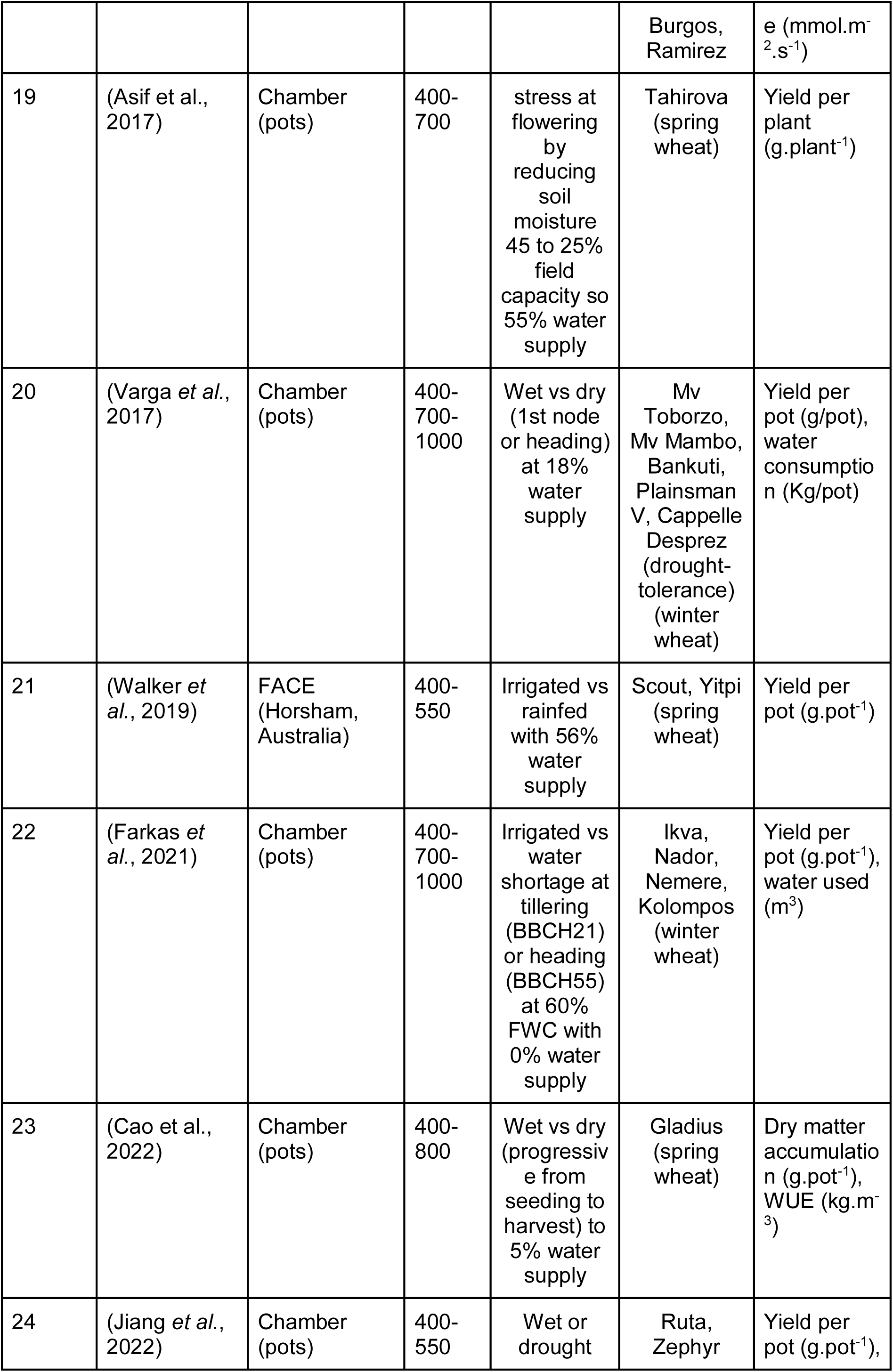
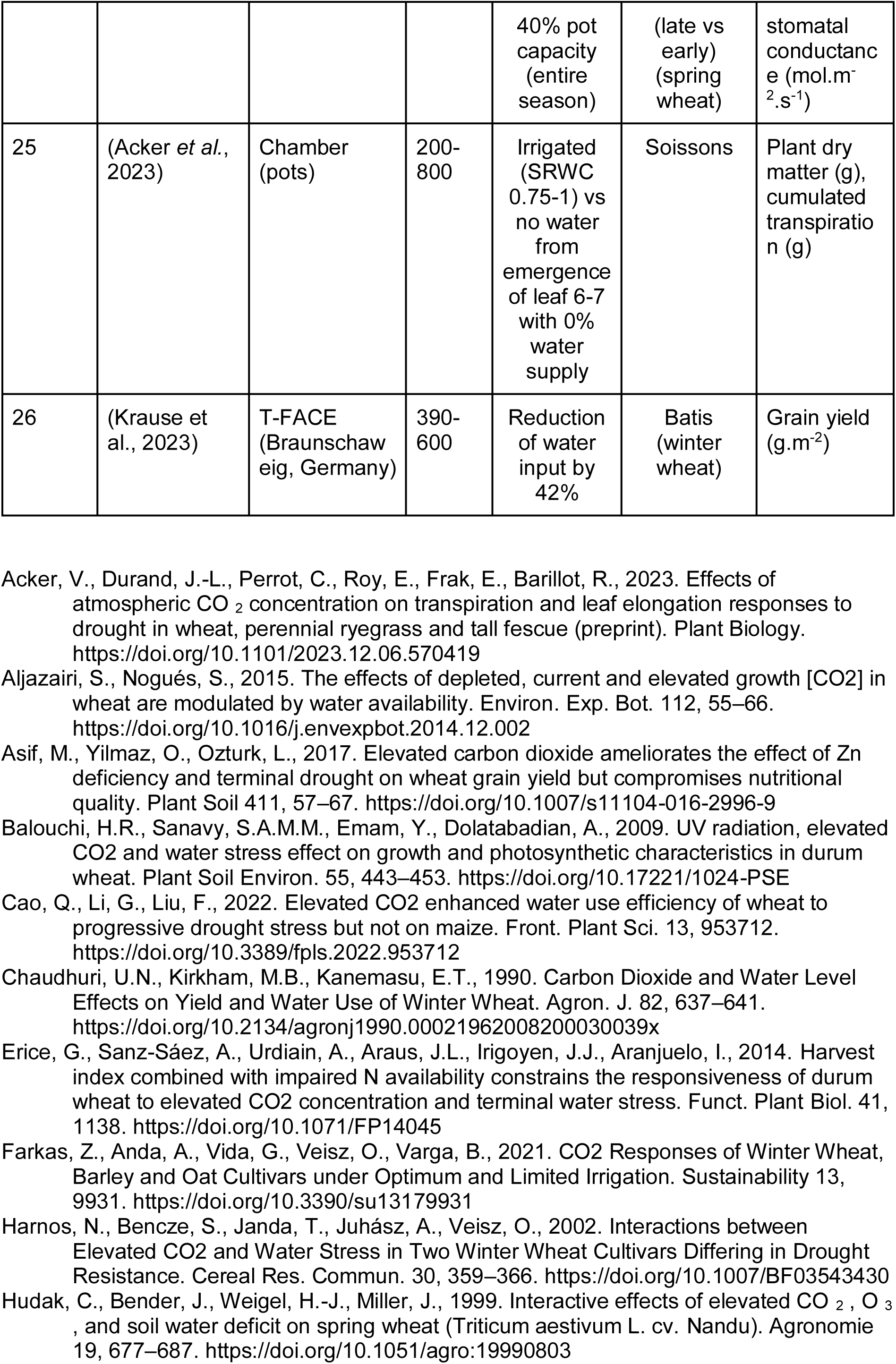
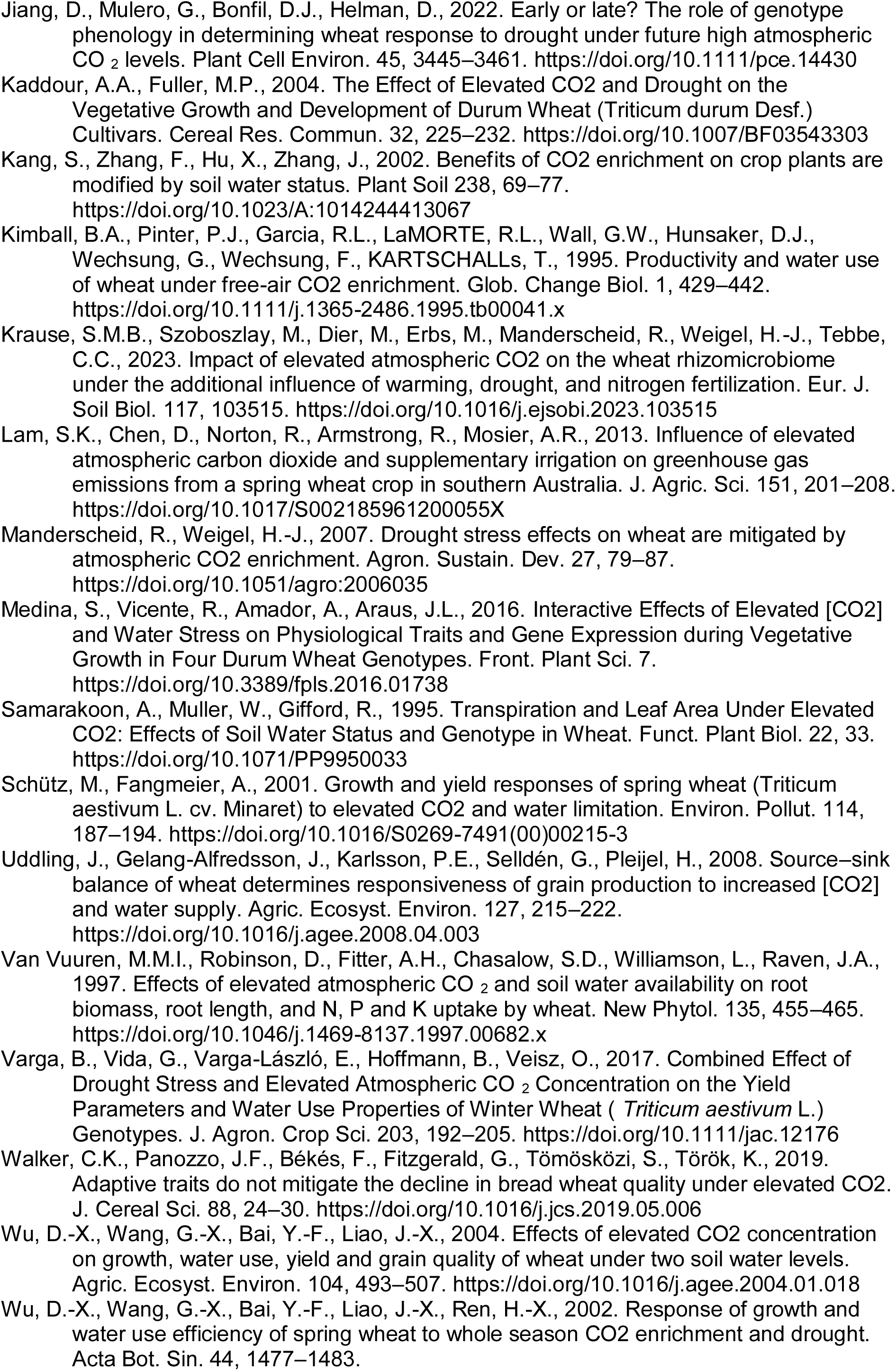
Characteristics from the studied references that reported results on crop productivity and/or water-related traits under elevated CO_2_ (eCO_2_) and water deficit (WD) conditions.

**Table S2.**
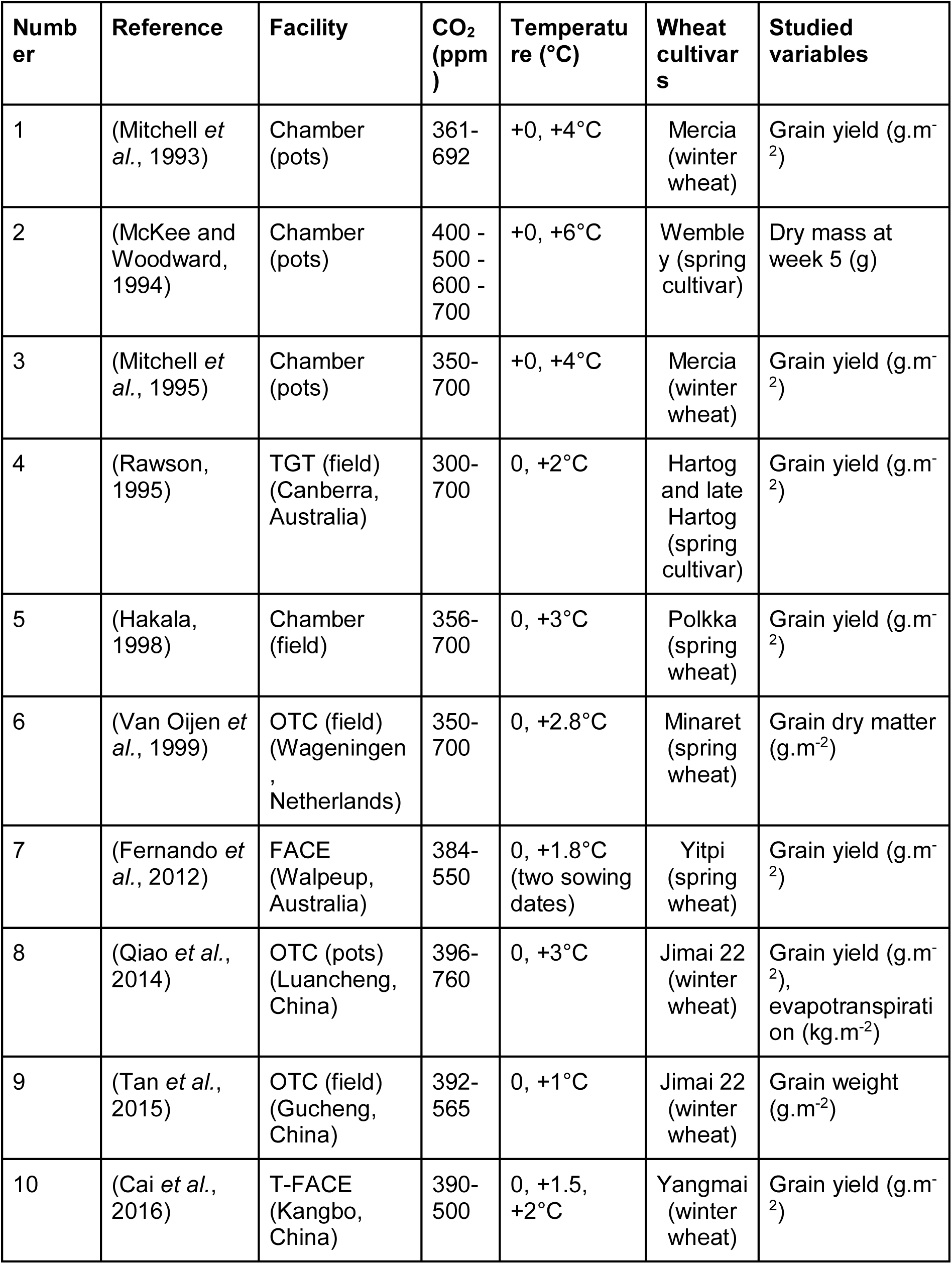
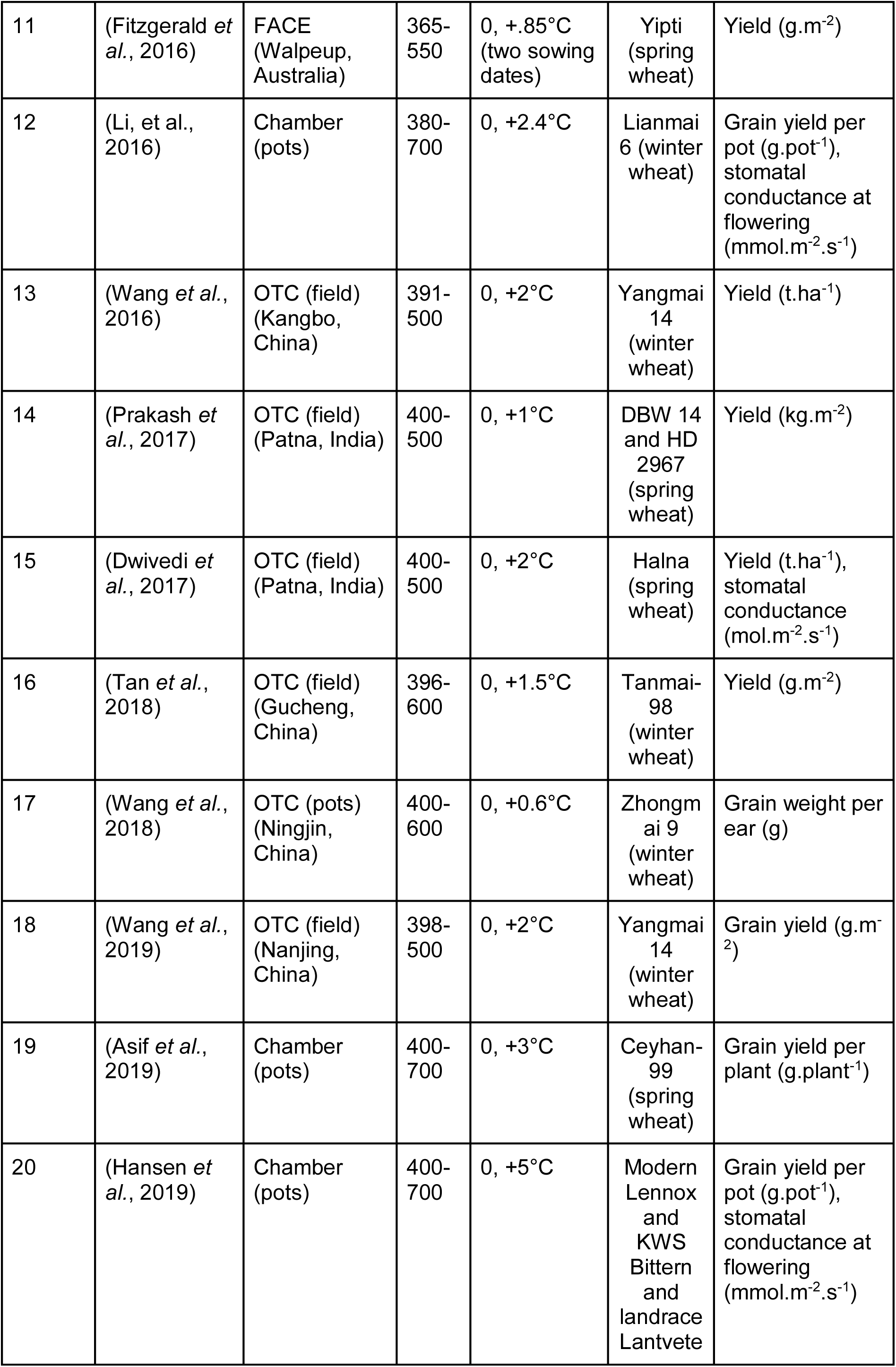
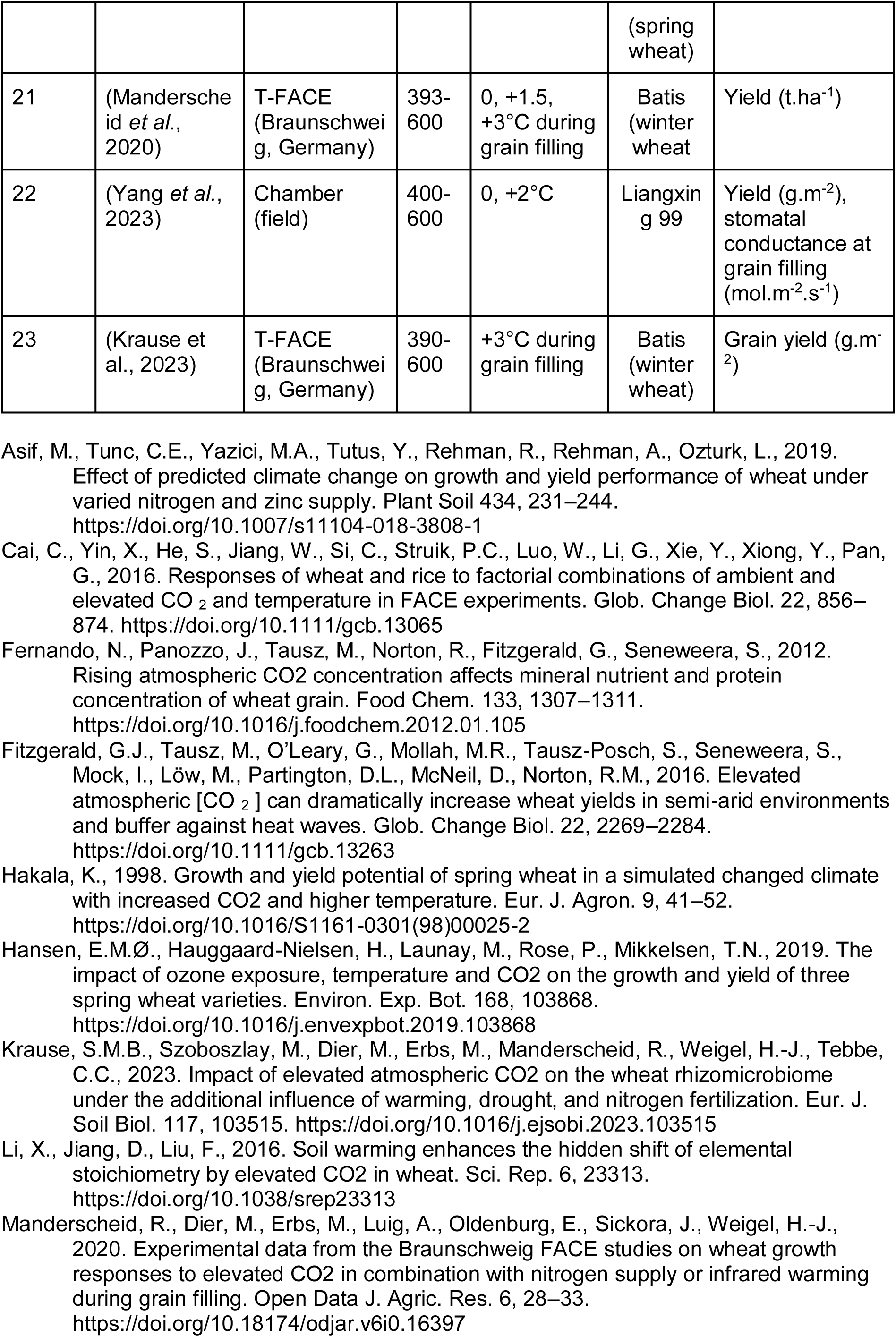
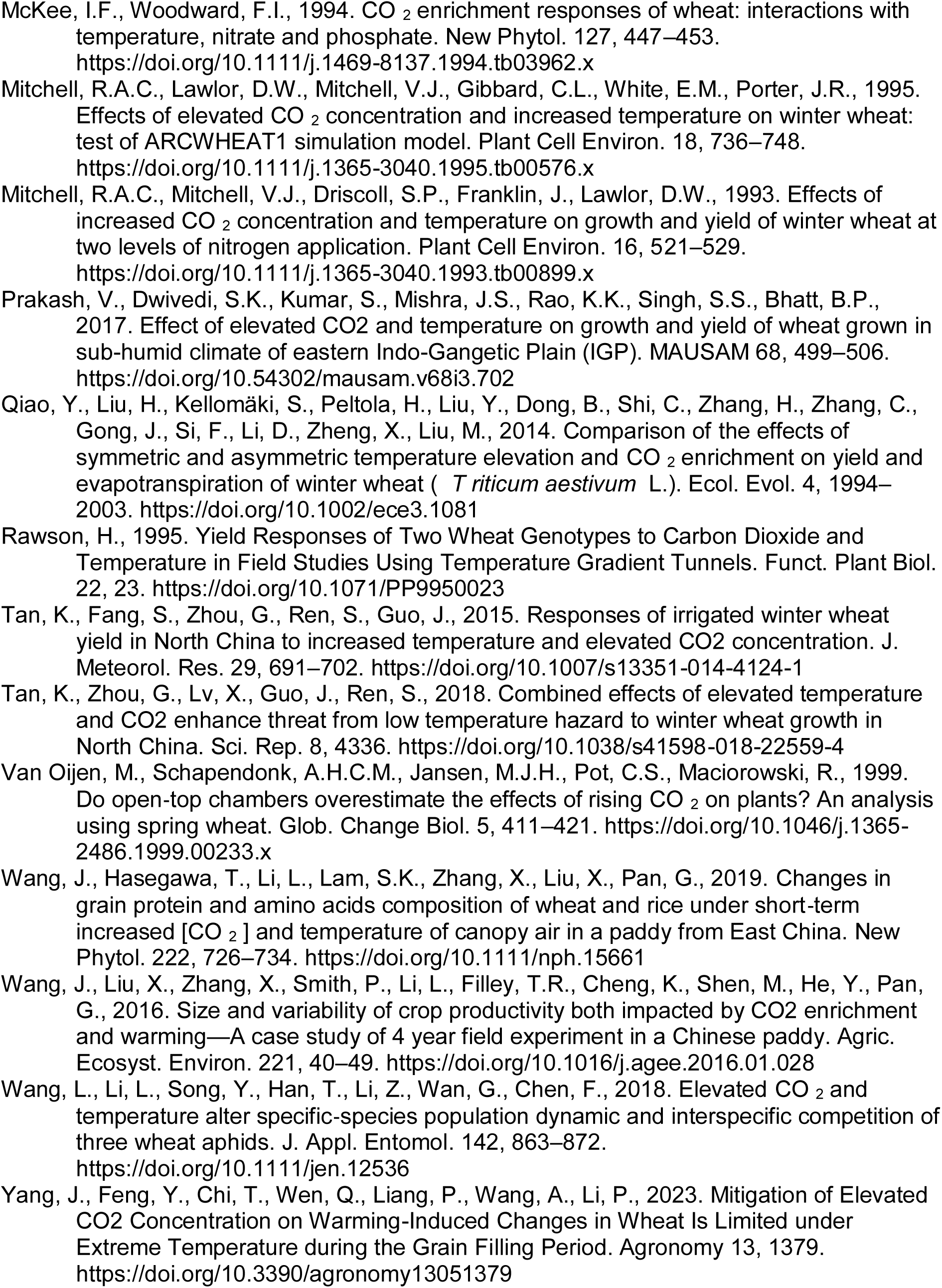
Summary of the extracted experimental studies with available data for crop productivity and water-related traits under eCO_2_ and high temperatures.

**Table S3.**
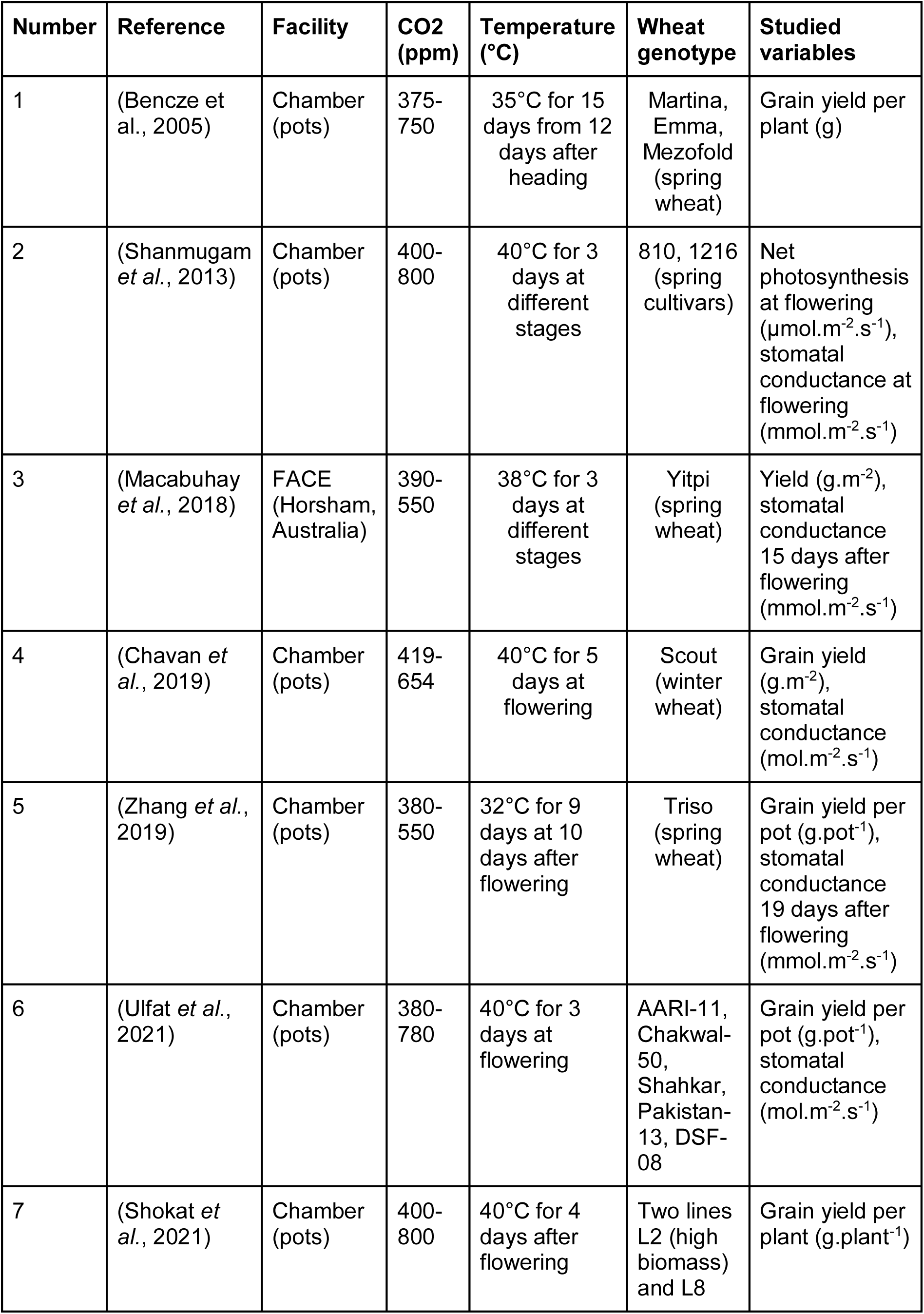
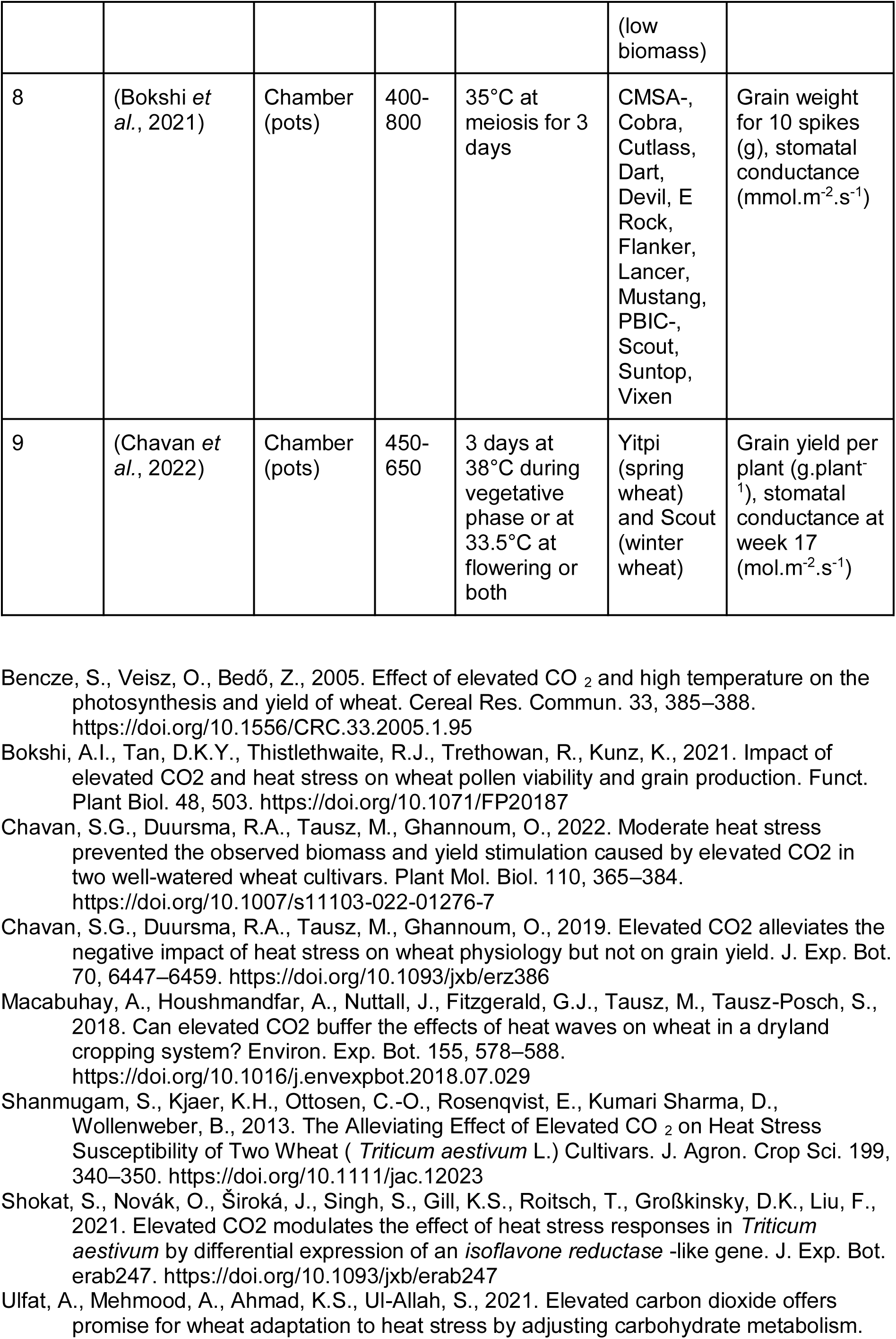
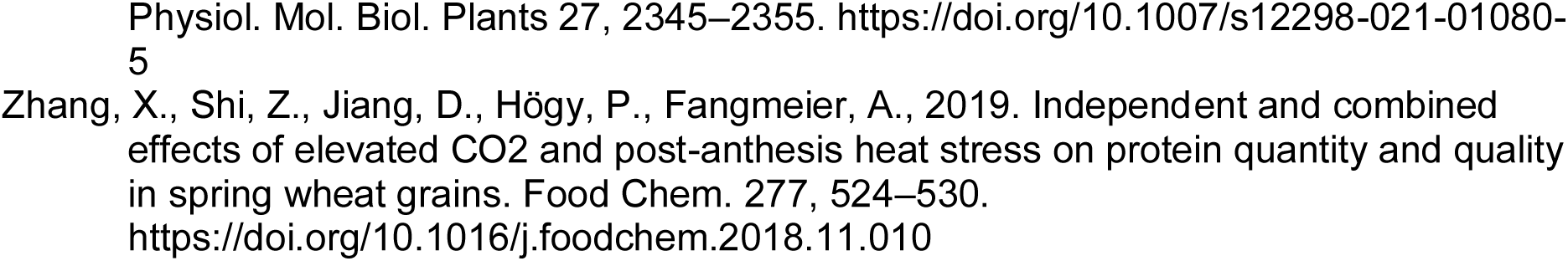
Summary of the extracted experimental studies with available data for crop performance and water-related traits under eCO_2_ and heat waves.

**Table S4.** Summary of the extracted experimental studies with available data for crop productivity and water-related traits under eCO_2_, water deficit and high temperature.

| Number | Reference | Facility | CO<sub>2</sub> (ppm) | Temperature (°C) | Drought | Interaction | Wheat genotype | Studied variables |
| --- | --- | --- | --- | --- | --- | --- | --- | --- |
| 1 | (Guoju *et al.*, 2005) | FACE (Haiyuan, China) | 360-450 | +0, +0.8, +1.8°C (different sowing dates) | With supplemental irrigation (30, 60, or 90mm) or without -> water supply at 0, 33, 67% | Concomitant | 79121-15 (spring wheat) | Yield (kg.hm<sup>-2</sup>) |
| 2 | (Dias De Oliveira *et al.*, 2013) | TGT (field) (Shenton, Australia) | 400-700 | +0, +2, +4, +6°C | Wet vs dry from flowering with 0% water supply | Concomitant | Janz non vigorous and 38-19 vigorous (spring wheat) | Yield (g.m<sup>-2</sup>) |
| 3 | (O’Leary *et al.*, 2015) | FACE (Horsham, Australia) | 365-550 | Two sowing dates (unspecified temperature difference) | Rainfed with 0% water supply vs irrigated | Concomitant | Yitpi | Grain yield (kg.ha<sup>-1</sup>), water used (mm) |
| 4 | (Dias De Oliveira *et al.*, 2015) | TGT (field) (Shenton, Australia) | 360-400 | +0, +3°C | Rainfed with 0% water supply vs irrigated | Concomitant | 7750N free tillering and 7750PF restricted tillering | Grain dry weight (mg), leaf transpiration rate (mmol.m<sup>-2</sup>.s<sup>-1</sup>) |
| 5 | (Fitzgerald *et al.*, 2016) | FACE (Horsham, Australia) | 365-550 | +0, +3°C (different sowing dates) | Wet vs dry with 77% water supply | Concomitant | Yipti, Janz (spring wheat) | Yield (g.m<sup>-2</sup>) |

**Table S5.** Summary of the extracted experimental studies with available data for crop productivity and water-related traits under eCO_2_, water deficit and heat waves.

| Number | Reference | Facility | CO<sub>2</sub> (ppm) | Temperature (°C) | Drought | Interaction | Wheat genotype | Studied variables |
| --- | --- | --- | --- | --- | --- | --- | --- | --- |
| 1 | (Li *et al.*, 2019) | Chamber (pots) | 400-800 | 40/35°C for 5 days at 5th day drought (stress) | Wet vs dry for 4 days after flowering with 61% water supply | Sequential | Gladius D-tolerant and Paragon D-sensitive (spring wheat) | Grain yield per plant (g.plant<sup>-1</sup>), stomatal conductance (mmol.m<sup>-2</sup>.s<sup>-1</sup>) |
| 2 | (Abdelhakim *et al.*, 2021) | Chamber (pots) | 400-800 | 36°C for 3 days at the end of tillering | Wet vs dry with 33% water supply | Concomitant | LM19, KU10 H-sensitive and LM62, GN5 H-tolerant (spring wheat) | Photosynthetic rate (μmol.m<sup>-2</sup>.s<sup>-1</sup>), stomatal conductance (mmol.m<sup>-2</sup>.s<sup>-1</sup>) |
| 3 | (Abdelhakim *et al.*, 2022) | Chamber (pots) | 400-800 | 36/28°C for 7 days at 2 days after sowing | Wet vs dry with 24% water supply | Heat stress, 3 days both, then drought stress | SF29 H-tolerant and LM20 H-sensitive (spring wheat) | Photosynthetic rate ($\mu\text{mol.m}^{-2}.\text{s}^{-1}$), stomatal conductance ($\text{mmol.m}^{-2}.\text{s}^{-1}$) |

- Abdelhakim, L.O.A., Mendanha, T., Palma, C.F.F., Vrobel, O., Štefelová, N., Čavar Zeljković, S., Tarkowski, P., De Diego, N., Wollenweber, B., Rosenqvist, E., Ottosen, C.-O., 2022. Elevated CO<sub>2</sub> Improves the Physiology but Not the Final Yield in Spring Wheat Genotypes Subjected to Heat and Drought Stress During Anthesis. *Front. Plant Sci.* 13, 824476. https://doi.org/10.3389/fpls.2022.824476 - Abdelhakim, L.O.A., Palma, C.F.F., Zhou, R., Wollenweber, B., Ottosen, C.-O., Rosenqvist, E., 2021. The effect of individual and combined drought and heat stress under elevated CO<sub>2</sub> on physiological responses in spring wheat genotypes. *Plant Physiol. Biochem.* 162, 301–314. https://doi.org/10.1016/j.plaphy.2021.02.015 - Li, C., Kuyper, T.W., Van Der Werf, W., Zhang, J., Li, H., Zhang, F., Hoffland, E., 2019. Testing for complementarity in phosphorus resource use by mixtures of crop species. *Plant Soil* 439, 163–177. https://doi.org/10.1007/s11104-018-3732-4

**Figure S1.**
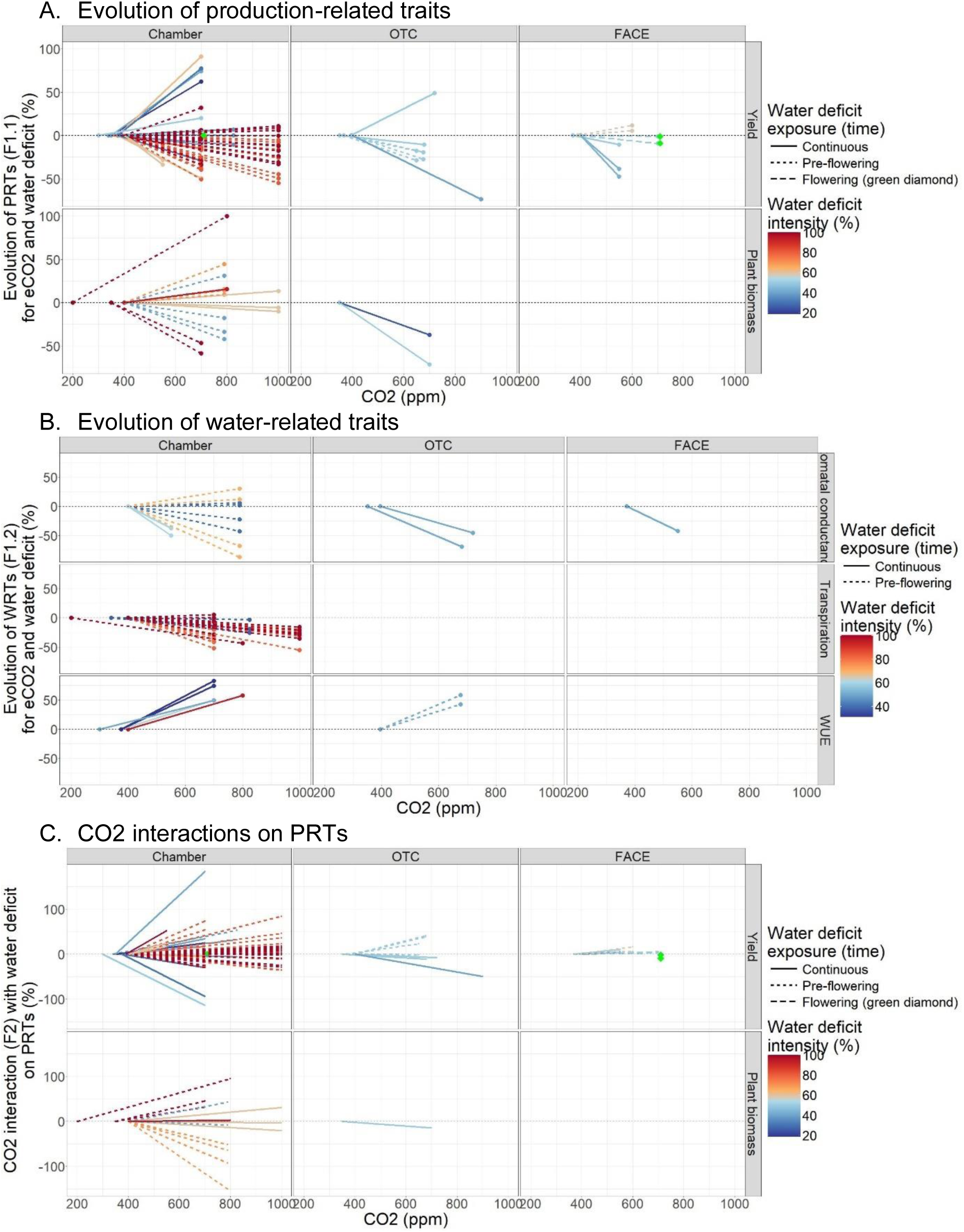
Relative reaction norms detailed with types of traits of plasticity indices for A) the evolution of production-related traits (PRTs) under future-like eCO2 and water deficit (F1.1, Δ_[aCO2_×_WW] vs [eCO2_×_WD]_ (PRT), Eq 1), B) the evolution of water-related traits (WRTs) under future-like eCO2 and water deficit (F1.2, Δ_[aCO2_×_WW] vs [eCO2_×_WD]_ (WRT), Eq 1) and C) the interaction between eCO2 and water deficit on PRTs (F2, Δ _eCO2, [WW] vs [WD]_ (PRT), Eq 2). Colors represent different water-deficit intensities (i.e. ratio of water inputs between treatments, in %) and line types represent different onset timings for soil water deficit. The black dashed horizontal lines indicate 0%. Relative reaction norms with stress onset at flowering were highlighted by green diamonds for better visibility.

**Figure S2.**
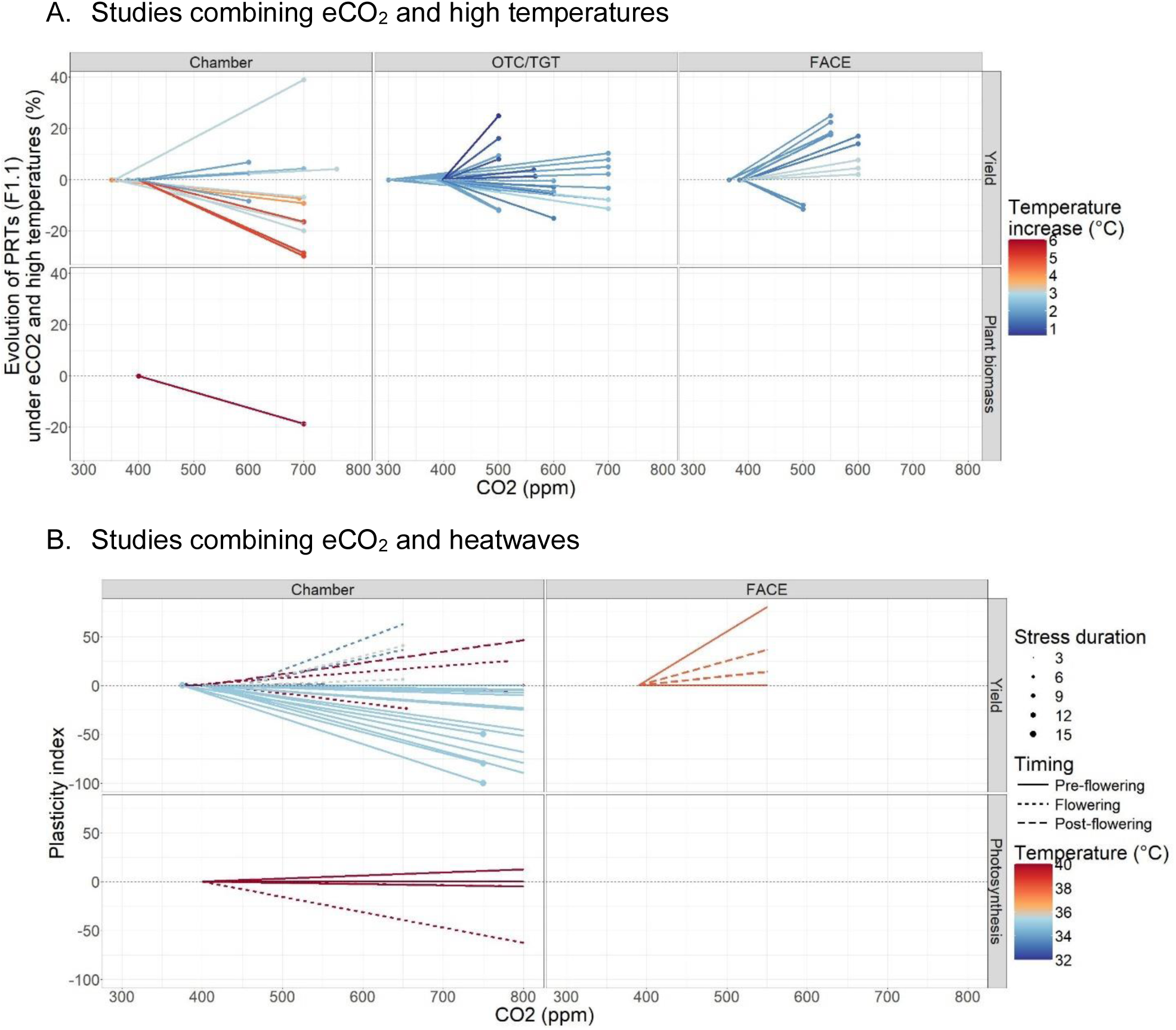
Relative reaction norms for productivity-related traits (PRTs) between non-stressed conditions under ambient CO_2_ (control) and treatments with eCO_2_ and warming applied as A) high temperatures throughout the growth cycle or the grain filling period with Δ _[aCO2 x aT] vs [eCO2 x HT]_ (PRT) and B) heatwaves with P Δ _[aCO2 x aT] vs [eCO2 x HW]_ (PRT) (Eq 1). Results for yield, above-ground plant biomass and photosynthesis presented for the different types of experiments, i.e. chambers, open-top chambers and temperature-gradient tunnels (OTC/TGT) and free-air CO_2_ enrichment (FACE). Colors represent different temperature increments for HT and stress temperature for heatwaves, line types represent different onset timings for heatwaves and point sizes represent the duration of heatwaves. The black dashed horizontal lines indicate 0%.

**Figure S3.**
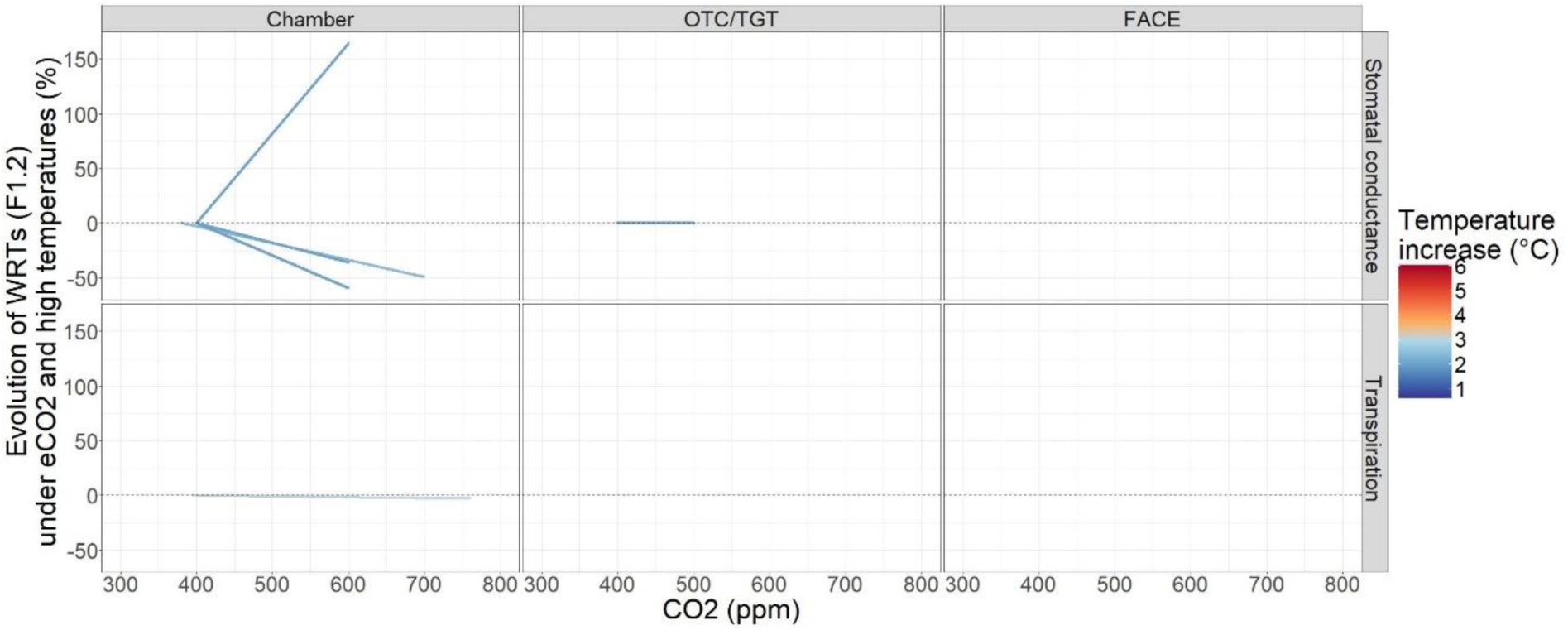
Relative reaction norms for stomatal conductance as a water-related trait (WRT) between ambient non-limiting conditions (control) and treatments with eCO_2_ and warming through high temperatures with Δ _[aCO2 x aT] vs [eCO2 x HT]_ (WRT) (Eq 1). Results presented for the different types of experiments, i.e. chambers and free-air CO_2_ enrichment (FACE). Colors represent stress temperature; line types represent different onset timings for heatwaves and point sizes represent the duration of heatwaves. The black dashed horizontal lines indicate 0%.

**Figure S4.**
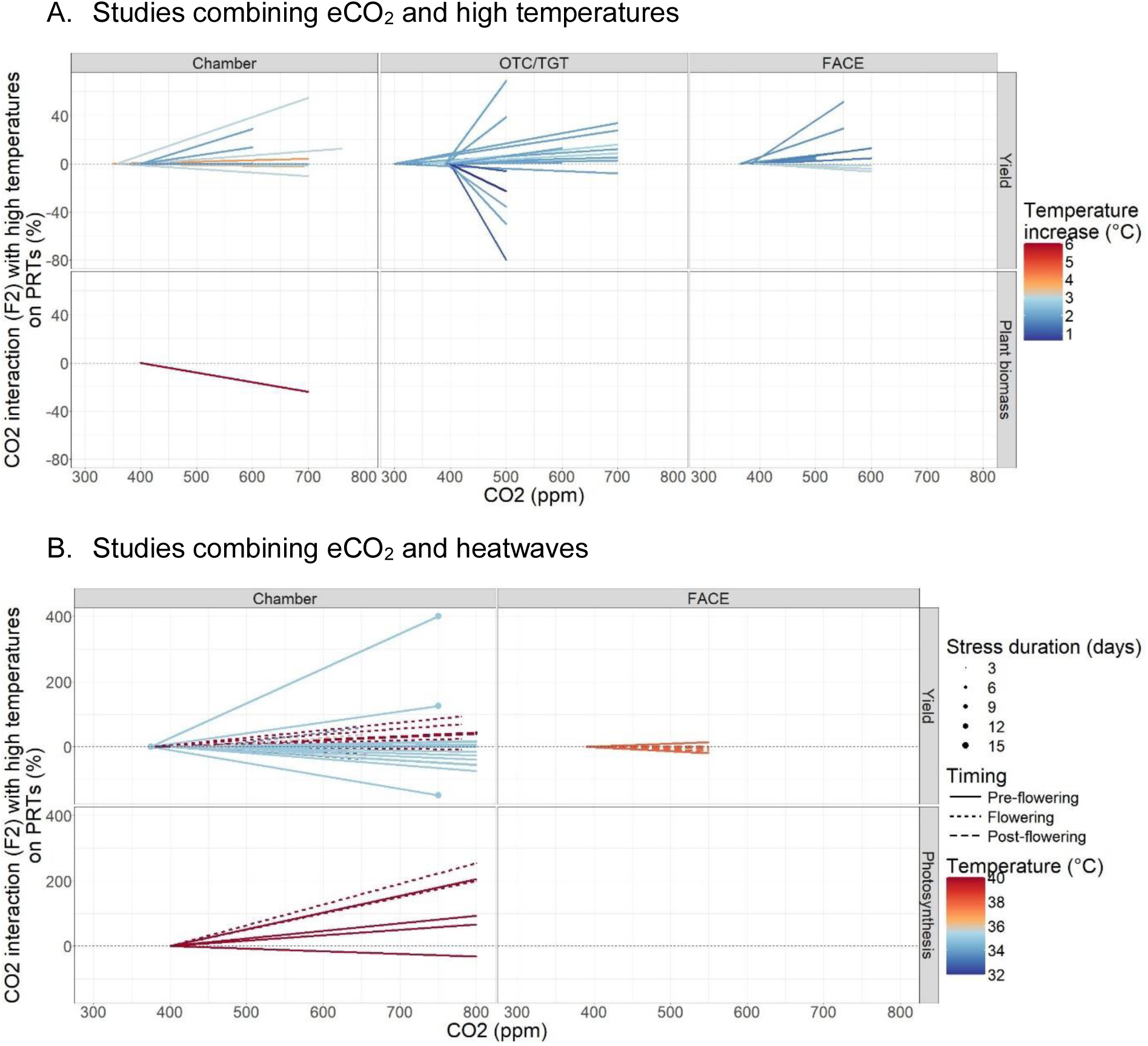
Relative reaction norms of plasticity indices for productivity-related traits (PRTs) between ambient non-limiting conditions (control) and treatments with eCO_2_ and warming through A) High temperatures with Δ _eCO2, [aT] vs [HT]_ (PRT) and B) Heatwaves with Δ _eCO2, [aT] vs [HW]_ (PRT) as the difference in relative eCO2 effects on PRTs at ambient temperature (aT) vs high temperature (HT) or heatwave (HW) (Eq 2). Results for yield and other variables presented for the different types of experiments, i.e. chambers, open-top chambers and temperature-gradient tunnels (OTC/TGT) and free-air CO_2_ enrichment (FACE) studies. Colors represent the temperature increment for HT and heatwave temperature for HW, line types represent different onset timings for heatwave and point size represent heatwave duration. The black dashed horizontal lines indicate 0%.

**Figure S5.**
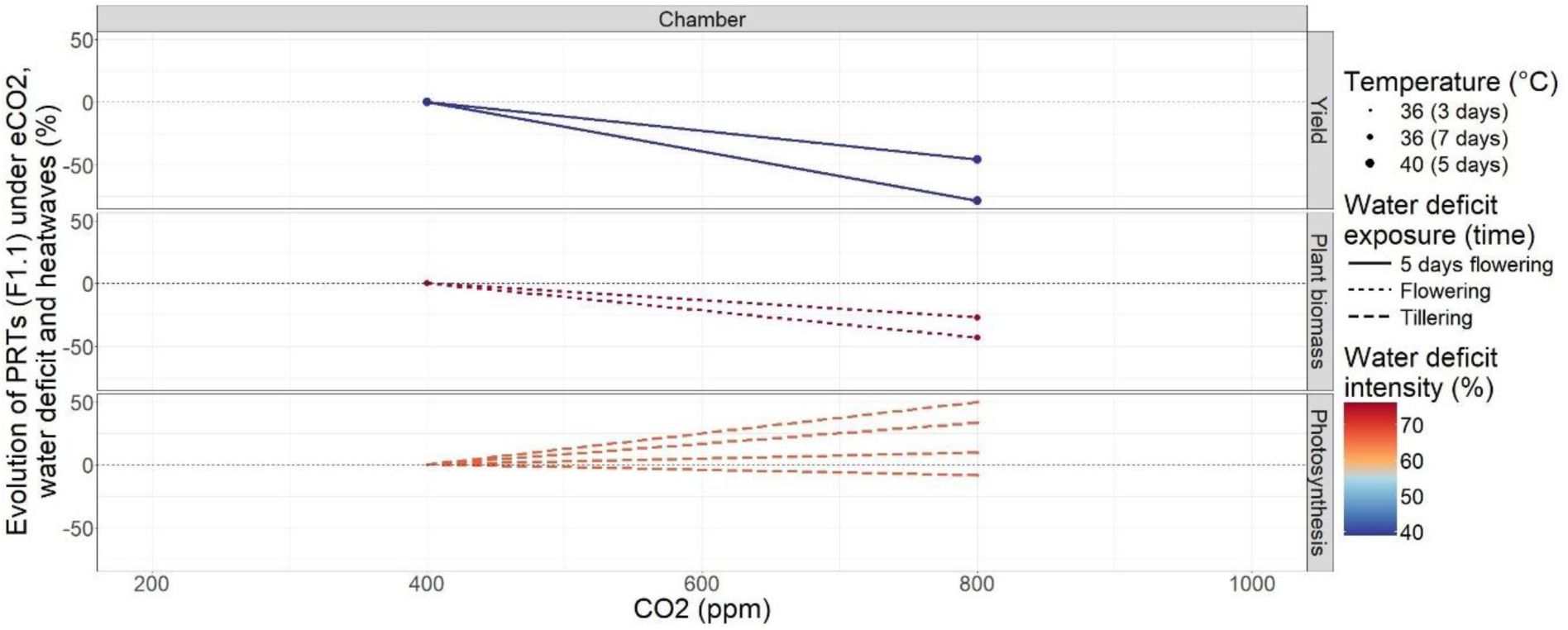
Relative reaction norms for productivity-related traits (PRTs) between ambient non-limiting conditions (control) and treatments with eCO_2_ and heatwaves with Δ _[aCO2 x WW x aT] vs [eCO2 x WD x HW]_ (PRT) (Eq1). Results for yield and other variables presented for open-top chamber/temperature gradient tunnel (OTC/TGT) and free-air CO_2_ enrichment (FACE) studies. Colors represent water deficit intensity (% based on water input), line types represent the type of water deficit applied and point sizes represent heatwave temperature and duration. The black dashed horizontal lines indicate 0%.

**Figure S6.**
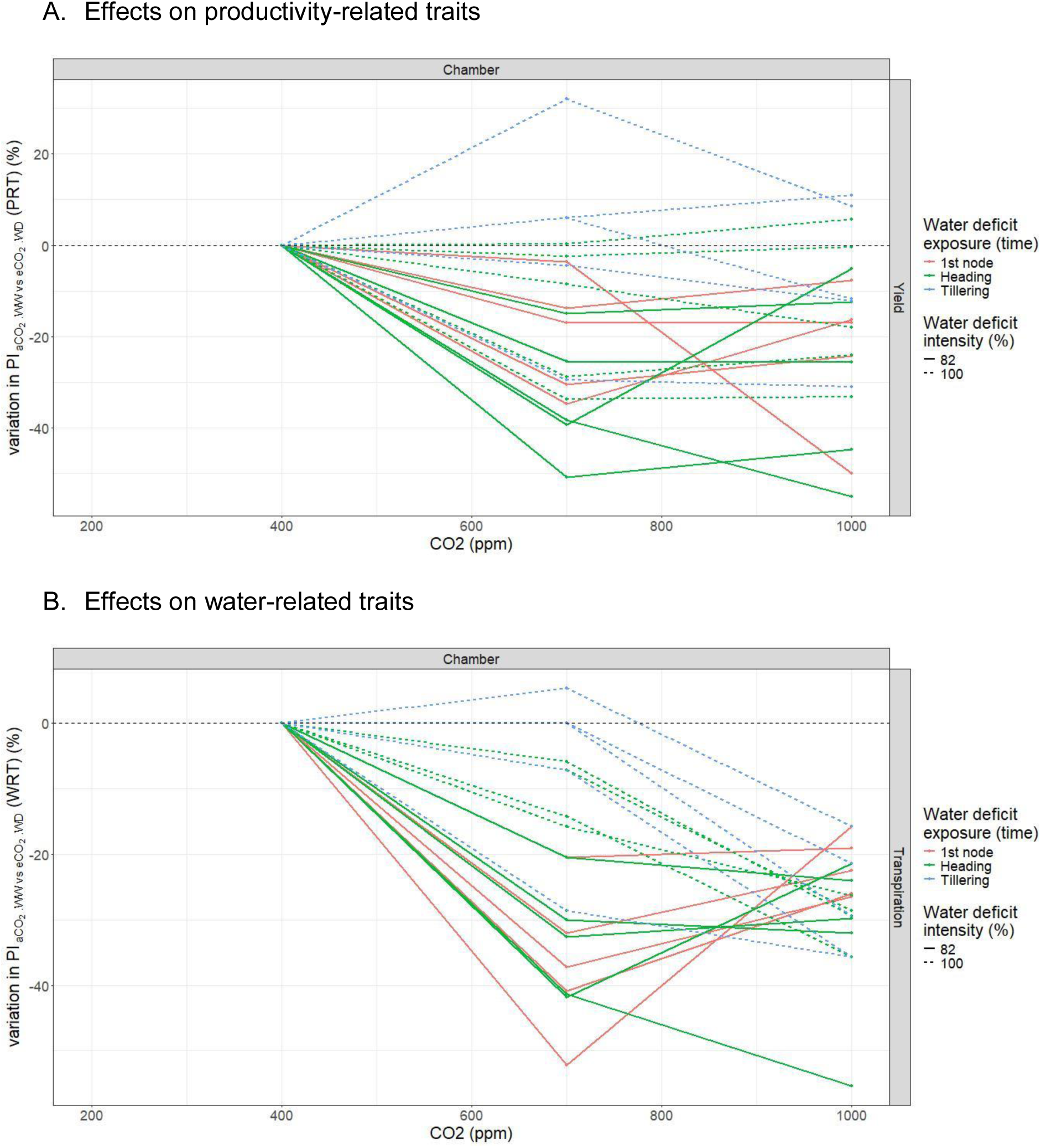
Relative reaction norms between non-stressed conditions with ambient CO_2_ (control) and treatments with different CO_2_ concentrations and water deficit for A) yield with Δ _[aCO2 x WW] vs [eCO2 x WD]_ (PRT) and B) Transpiration with Δ _[aCO2 x WW] vs [eCO2 x WD]_ (WRT). Line types represent water deficit intensity (% based on water input), colors represent the type of water deficit applied. The black dashed horizontal lines indicate 0%. Yield response shows a saturating effect between 700 and 1000 ppm in most cases but with some exceptions. For transpiration, relative reactions norms were mostly linear for data from Farkas et al. (2021) but saturating or with lower decrease at 1000 pm in data from Varga et al. (2017).

